# A Synthetic Microbial Therapy Rewires Antitumor Immunity Across Multiple Cancer Types

**DOI:** 10.64898/2026.07.19.735925

**Authors:** Jugal K. Das, Qing-Ming Qin, Shailbala Singh, Christina James Thomas, Shreyan Gupta, Ayan Chatterjee, Sunilgowda Sunnagatta Nagaraja, Fengguang Guo, Kaylee Delgado, Anil Kumar, Esther Ryu, Bennett Flannagan, Cansu Agca, Yuksel Agca, Noah Powell, Erin Barry, Melissa M. Kahl-McDonagh, Sankar P. Chaki, Andre Mendes Ribeiro Correa, Seyednami Niyakan, Song-I Han, James J. Cai, Xiaoning Qian, Koichi S Kobayashi, Arum Han, Arul Jayaraman, Thomas A. Ficht, Leslie Garry Adams, Robert C. Alaniz, Cassian Yee, Jianxun Song, Paul de Figueiredo

## Abstract

Microbial immunotherapies show promise against cancer, yet broad efficacy and mechanistic insight remain elusive. Here, we introduce SPIKE 1.0 (S1.0), a metabolically engineered bacterium that converts tryptophan into immunomodulatory hydroxyindoles via tryptophan monooxygenase to remodel the tumor microenvironment (TME). A single systemic dose of S1.0 elicited potent, durable antitumor responses across multiple murine models, including humanized mice, with minimal toxicity. S1.0 enhanced inflammatory signaling, activated innate and adaptive immunity, and promoted T cell persistence, memory, and resistance to exhaustion. It outperformed checkpoint inhibitors and synergized with chemotherapy. Multi-omics profiling revealed that S1.0 rewired amino acid metabolism in tumor-infiltrating immune cells and disrupted immunosuppressive networks. These results establish S1.0 as a scalable, cost-effective microbial immunotherapy with broad translational potential for solid tumors.

---

Bacteria-based cancer immunotherapy (BCIT) has emerged as a transformative strategy for cancer treatment, offering unique advantages over conventional therapies^1,2^. Therapeutic bacteria preferentially accumulate and proliferate within the tumor microenvironment (TME), disrupting tumor cell metabolism, enhancing antitumor immune responses and reversing immune suppression. Furthermore, therapeutic bacteria can precisely deliver therapeutic agents, including cytokines, antibodies, and other anticancer molecules directly to tumors. BCIT has shown promise as both a monotherapy and in combination with other modalities, improving clinical outcomes^3,4^. However, achieving consistent and effective cancer regression across diverse cancer types by using BCIT remains a major challenge, and the molecular mechanisms driving bacterial efficacy remain poorly understood.

*Brucella melitensis* 16MΔ*vjbR* (BmΔ*vjbR*) is a genetically attenuated strain lacking the virulence regulator VjbR, a LuxR-type transcription factor that controls expression of the virB operon and other Type IV secretion system (T4SS)–associated virulence genes required for intracellular survival and establishment of infection^5^. Our recent work demonstrated that the live attenuated strain of Bm*ΔvjbR* is a promising chassis for therapeutic development^6^. This strain exhibited minimal endotoxin activity, genetic tractability, and a strong safety profile across various immune-sufficient and –deficient mouse models^7–11^. Bm*ΔvjbR* specifically homed to tumor tissues, remodeled the TME, promoted proinflammatory M1 macrophage polarization, and enhanced CD8^+^ T cell infiltration and activity. This strain synergized with adoptive cell transfer (ACT) therapies^6^, highlighting its promise as an adjunct to immunotherapy. However, Bm*ΔvjbR* showed limited efficacy across tumor types, particularly displaying sub-optimal activity against melanoma. We therefore engineered BmΔ*vjbR* to produce immunostimulatory tryptophan-derived metabolites, with the goal of enhancing and broadening its antitumor effects while uncovering the molecular mechanisms driving improved activity against solid tumors.

Tryptophan-derived metabolites, such as indole and its derivatives, modulate T cell fate and function^12,13^. We have previously shown that indole promotes regulatory T cell (Treg) activity and reduces inflammation when produced by engineered BmΔ*vjbR* in a rheumatoid arthritis model^14^. Related metabolites, including indole-3-propionic acid (IPA), enhance CD8⁺ T cell–mediated responses in pancreatic cancer, with other indole derivatives under investigation for anticancer potential^15,16^. However, their role in immunomodulation remains largely unexplored. Emerging evidence highlights the critical role of the microbiome in reshaping the immune landscape of the TME, particularly by regulating the metabolic precursors essential for T cell function, such as glutamine and tryptophan^17,18^. We hypothesized that engineering Bm*ΔvjbR* to produce indole derivatives would enhance and broaden its anti-cancer efficacy and sought to define the cellular and molecular mechanisms driving efficacy among diverse tumor models.

We developed SPIKE 1.0 (Synthetic Programmable bacteria for Immune-directed Killing in tumor Environments), a metabolically engineered strain that constitutively produces hydroxyindoles (HIs) to reprogram the suppressive TME. SPIKE 1.0 enhanced tumor immunogenicity, restored exhausted T cell function, and demonstrated greater efficacy across multiple refractory cancers. By remodeling amino acid (AA) bioavailability in intratumoral T cells, it sustained potent antitumor immunity. This work establishes SPIKE 1.0 as a scalable, cost-effective platform that integrates metabolic reprogramming with immunotherapy to overcome resistance in solid tumors.

## Results

### HI boosts CD8+ T cell killing ability and increases proinflammatory CD4+ T cell activity

Some indole derivatives, including HI derivatives, display immunomodulatory activities^19,20^. However, our understanding of the specific ways in which indole and its derivatives alter immune responses remains incomplete. We therefore tested the hypothesis that HIs exhibit potent immunomodulatory effects. Our findings demonstrated that 5-HI augmented Th1 and Th17 responses in CD4^+^ T-cells, thereby driving antitumor immunity by releasing the cytokine INF-γ (Figure S1A). 5-HI also significantly promoted several anticancer activities in immune cells and therefore was pursued further in this work. Specifically, we determined by flow-cytometric analysis using the illustrated gating strategy (Figure S1B-D) that 5-HI increased the production of granzyme B (GrB), TNF-α, and perforin, in murine CD8^+^ T cells (Figure 1A-B). This metabolite also promoted the differentiation of murine CD4^+^ T cells into a Th17 inflammatory subtypes (Figure 1C-D) and the polarization of macrophages towards an M1-like phenotype (Figure 1E-F), as indicated by increased CD38 expression in murine bone marrow-derived macrophages (BMDMs). Moreover, 5-HI significantly enhanced TNF-α production in human CD8^+^ T cells (Figure S1E-F) and murine CFSE^+^ (carboxyfluorescein succinimidyl ester) CD8^+^ T cells (Figure S1G-H), but not in murine CD4^+^ T cells (Figure S1I-J), thereby extending findings from murine to human cells.

**Figure 1.**
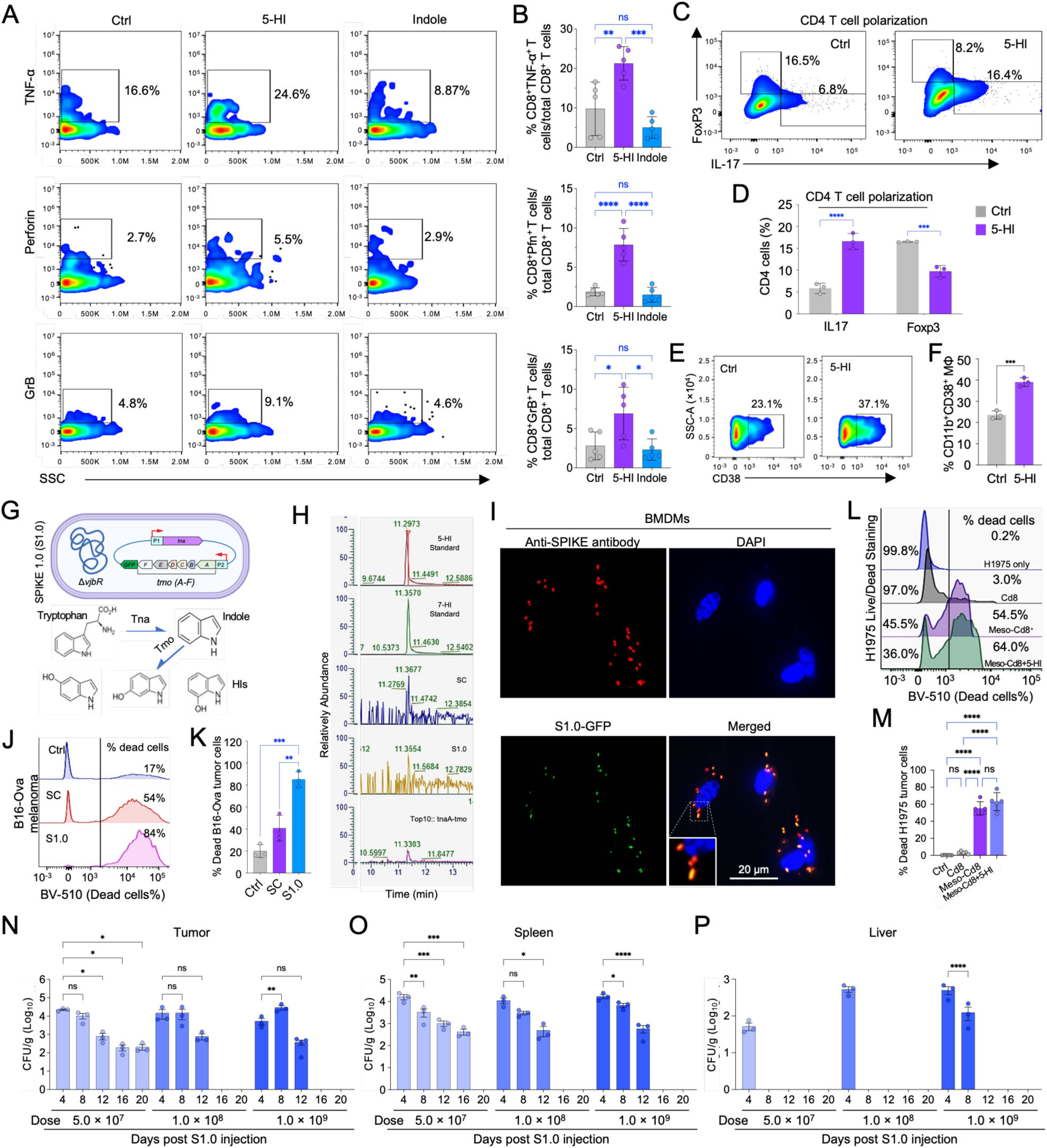
Hydroxyindole (HI) activates CD8^+^ T cells and enhances their cytotoxic activity. (A) Representative flowcytometric analysis of proinflammatory cytokine TNF-α and cytotoxic markers perforin, granzyme B (GrB) in mouse CD8^+^ T cells pooled from spleen and lymph nodes of mice from a single group. Ctrl: control. (B) Quantitative analysis of flowcytometric dot-plot plots represented in panel (A). (C, D) CD4^+^ T cells differentiation into Th17/Treg cell types upon treatment with HI. Representative flowcytometric analysis (C) and quantitative analysis of CD4^+^ T cells differentiation into T helper type 17 (Th17) cells or T regulatory (T_reg_) cells upon HI treatment (D). (E) Representative flowcytometric assessment of M1 macrophage polarization by CD38 expression analysis. (F) Quantitative of CD11b^+^CD38^+^ M1 macrophage polarization represented in panel (E). (G) Illustration of the genetic circuit used to construct SPIKE 1.0 (S1.0) bacterial strain producing hydroxyindole. (H) Detection of HIs produced by the bacterial S1.0 determined by qualitative LC-MS/MS analysis. (I) Immunofluorescence analysis of the intracellular S1.0 in mouse bone marrow-derived macrophages (BMDMs). (J) Representative dead-live flowcytometric assay (DLFA) of B16-Ova melanoma cells co-cultured with OT-1 CD8^+^ T cells that had been pre-cocultured with S1.0, SPIKE-chassis (SC), or 1× PBS control (Ctrl) treated bone marrow-derived macrophages (BMDMs). (K) Quantitation of dead B16-Ova cells represented in panel (J). (L-M) Representative DLFA of mesothelin antigen specific CD8^+^ T cells co-cultured with H1975 lung cancer cells (L) and quantitation of dead H1975 cells in the indicated treatments (M). The representative DLFA data (J-M) is shown from a CD8^+^ T cells to cancer cells ratio of 1:1. (N-P) S1.0 distribution in tumor (N), spleen (O), and liver (P) in Lewis lung carcinoma (LLC1) tumor-bearing mice administrated with various doses of S1.0. n = 10/group. Quantitative data represent mean ± standard error of measurement (SEM) from three independent experiments. *, **, ***, ****, *p* < 0.05, 0.01, 0.001, and 0.0001, respectively.

We hypothesized that the pro-inflammatory properties of HIs could be harnessed to modulate the TME and enhance antitumor immunity in solid tumors. To test this possibility, we first genetically engineered the anticancer strain BmΔ*vjbR* (SPIKE chassis, hereinafter SC)^6^ to express tryptophanase (Tna) and toluene-4 monooxygenase (Tmo) genes, generating SPIKE 1.0 (S1.0), which constitutively produced HIs (Figure 1G-H). Fluorescence microscopy confirmed the production of HIs did not impair host cell uptake, as S1.0 entered and persisted within BMDMs (Figure 1I). Next, we tested the hypothesis that S1.0 displayed enhanced antitumor activities and found that CD8^+^ T cells co-cultured with S1.0 induced significantly higher B16-Ova melanoma tumor cell killing than the SC parental strain, and that S1.0 had significantly higher cytotoxic activity compared to SC as demonstrated by flow-cytometric analysis (Figure 1J-K). Finally, treatment with 5-HI also significantly enhanced the cytotoxic potential of human mesothelin (Meso)-specific CD8^+^ T cells against New York esophageal squamous cell carcinoma 1 (NY-ESO-1)-expressing H1975 human lung tumor cells, and murine meso-Panc02 pancreatic cancer cells (Figure 1L-M, Figure S1K-L). Marker panels for murine and human CD8⁺ T cells were optimized independently to account for species-specific differences in antibody performance and functional readouts. Murine CD8⁺ T cells were assessed using GrB, perforin, and TNFα as established indicators of cytotoxic effector function. In contrast, human CD8⁺ T cells were evaluated using anti-inflammatory TNFα and IL-2, which are validated markers of activation, functional competence, and exhaustion *in ex vivo* human T cell assays^21^. This approach ensured biologically relevant and technically robust assessment of CD8⁺ T cell responses in each species. Taken together, our results demonstrated that the pro-inflammatory activity of HI induced cytotoxic activity against diverse types of cancer cells.

The *in vitro* anti-tumor efficacy of S1.0 encouraged us to examine its safety profile in multiple animal models. In the mouse model, organ weights and complete blood count (CBC) analyses showed no significant differences in mice treated with either 1×10^7^ or 1×10^8^ CFU (colony forming unit) of S1.0, doses mimicking our intervention dosages in solid tumors (Figure S2A-B). S1.0 persisted in tumor and spleen for 12 days post-injection (DPI). However, it could be recovered only from tumor tissue and no other organs in tumor-bearing mice at 20 DPI (Figure 1N-P, Figure S2C-F). When tumor-bearing mice were treated with a combination of S1.0 and adoptively transferred CD8^+^ T cells, viable S1.0 was not detected in any of the tested organs of treated mice at 12 DPI (Figure S2D), implying complete tumor regression in the treated mice. In a pregnant goat model, a sensitive testbed for evaluating adverse events caused by bacterial therapeutic candidates, no abortions in mid-gestational animals were detected following S1.0 treatment (Figure S2G). Also, we failed to recover any viable S1.0 from specimens of dams throughout the experiments (Figure S2G). Collectively, these data demonstrated the favorable safety features of S1.0 in these tested animal systems.

### S1.0 inhibits tumor growth and improves survival in multiple murine tumor models

To test whether S1.0 displays anticancer activity in multiple mouse models of solid tumors, we evaluated both tumor growth kinetics and the survival of tumor-bearing mice treated with a single intravenous dose of the intervention. We used two syngeneic heterotopic murine cancer models B16-Ova melanoma (warm tumor) and pancreatic cancer Panc02 (cold tumor) (Figure 2A). To extend our findings to human solid tumor models, we also used lung cancer (H1975) and pancreatic cancer (AsPC-1) heterotopic xenograft models (Figure 2B), in which immunocompromised non-obese diabetic (NOD)-scid IL2rγ^null^ (NSG) mice were engrafted with these tumor cells. The NSG mouse model is well-suited for evaluating CAR-T cell efficacy, as its deficiency in endogenous immune cells, including B, T, and natural killer (NK) cells, minimizes confounding adaptive immune responses^22^. Moreover, because NSG mice do not reject human tissues, they allow for the accurate evaluation of how S1.0 immunomodulates the TME of human tumors and human immune cells within a living system. Results from these experiments showed that in both syngeneic (Figure 2C-F) and xenograft (Figure 2G-J) mouse-tumor models, S1.0 treatment significantly reduced tumor sizes and increased the survival of tumor (i.e., B16-Ova melanoma, Panc02, H1975, and AsPC-1)-bearing mice (Figure 2C-J). S1.0 significantly out-performed the parental SC in anti-cancer treatment efficacy in both mouse tumor models (B16-Ova melanoma and Panc02) and human solid tumor models of H1975 and AsPC-1 tumor (Figure 2C-J). Next, we extended our studies to a KPC-luciferase/Luc orthotopic pancreatic ductal adenocarcinoma (PDAC) syngeneic mouse model^23^ (Figure 3A) and found that a single dose treatment with S1.0 inhibited PDAC tumor growth, and extended survival to a similar level as treatment with 6 doses of immune checkpoint inhibitor (ICI) αPD-1 (anti-PD-1) antibody; moreover, the survival benefit was further enhanced (two-fold) when mice were treated with a combination of S1.0 and αPD-1 therapy (Figure 3B-E). Importantly, in some animals, treatment with S1.0 alone or combined with αPD-1 led to complete resolution of orthotopic pancreatic tumors, with no recurrence observed until the experiment ended (Figure 3B-E), suggesting that a single dose of S1.0 contributed potent anticancer activities.

**Figure 2.**
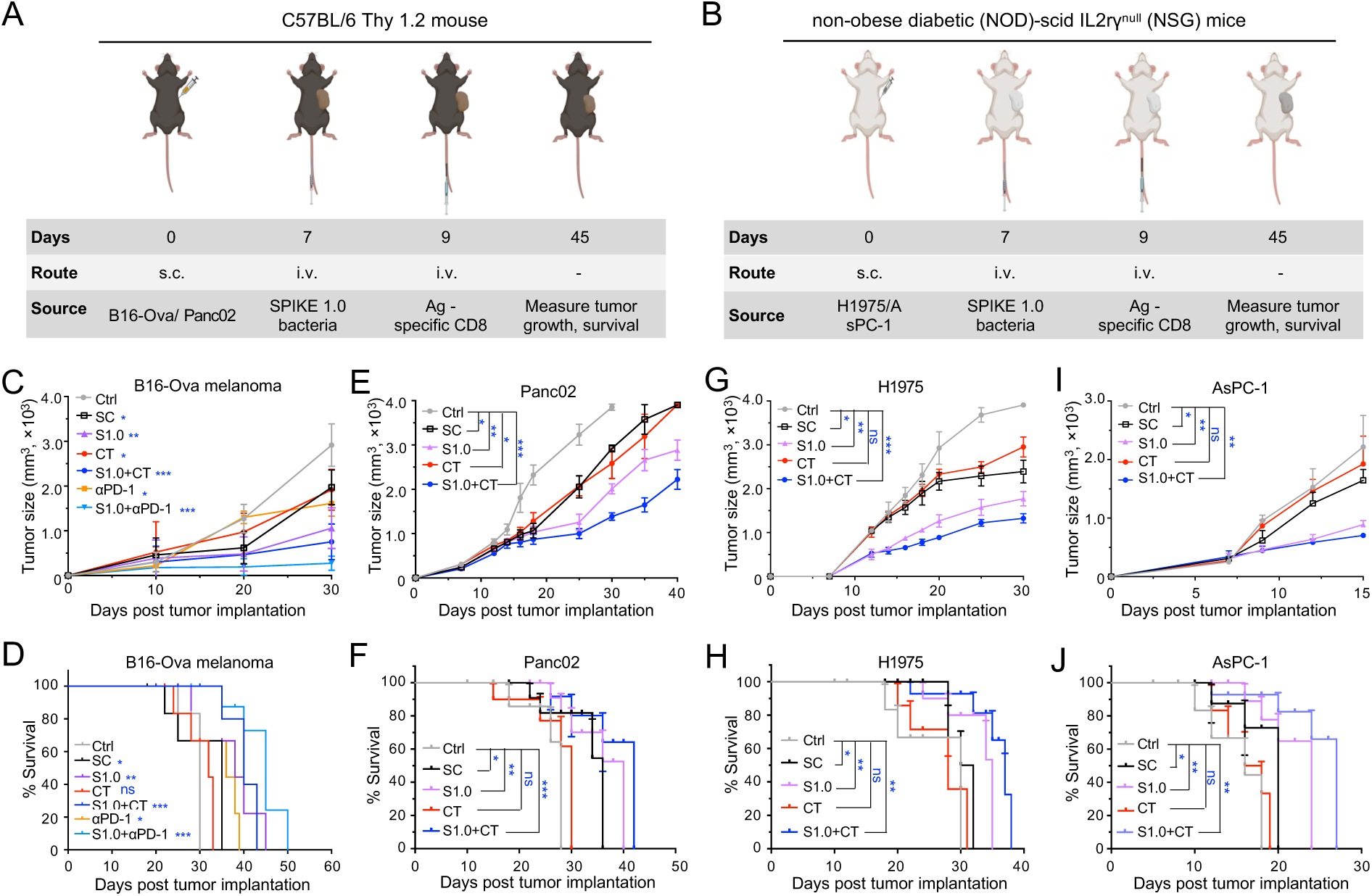
S1.0 inhibits tumor growth and increases survival of tumor-bearing mice. (A, B) Schematic illustration of mouse tumor model experiments in B6 wildtype mice (B16-Ova melanoma and Panc02 tumors) (A) and in NSG mice (H1975 and AsPC-1 tumors) (B). s.c: subcutaneous, i.v: intravenous. (C-F) Tumor growth and survival curves of C57BL/6 mice bearing B16-Ova melanoma (C, D) and Panc-02 (E, F) tumors. Ctrl, SC, S1.0, CT, αPD-1: control, SPIKE chassis, SPIKE 1.0, CAR T-cell transfer/therapy, and anti-PD-1 antibody, respectively. (G-J) Tumor growth curves of human H1975 lung tumor (G) and mice survival curves (H) and human pancreatic AsPC-1 tumor growth (I) and mice survival curve (J) in immunocompromised nonobese diabetic/severe combined immunodeficiency (NOD/SCID) γ mice. *, **, ***, and ****: p ≤ 0.05, 0.01, 0.001, and 0.0001, respectively. ns. Not significant.

**Figure 3.**
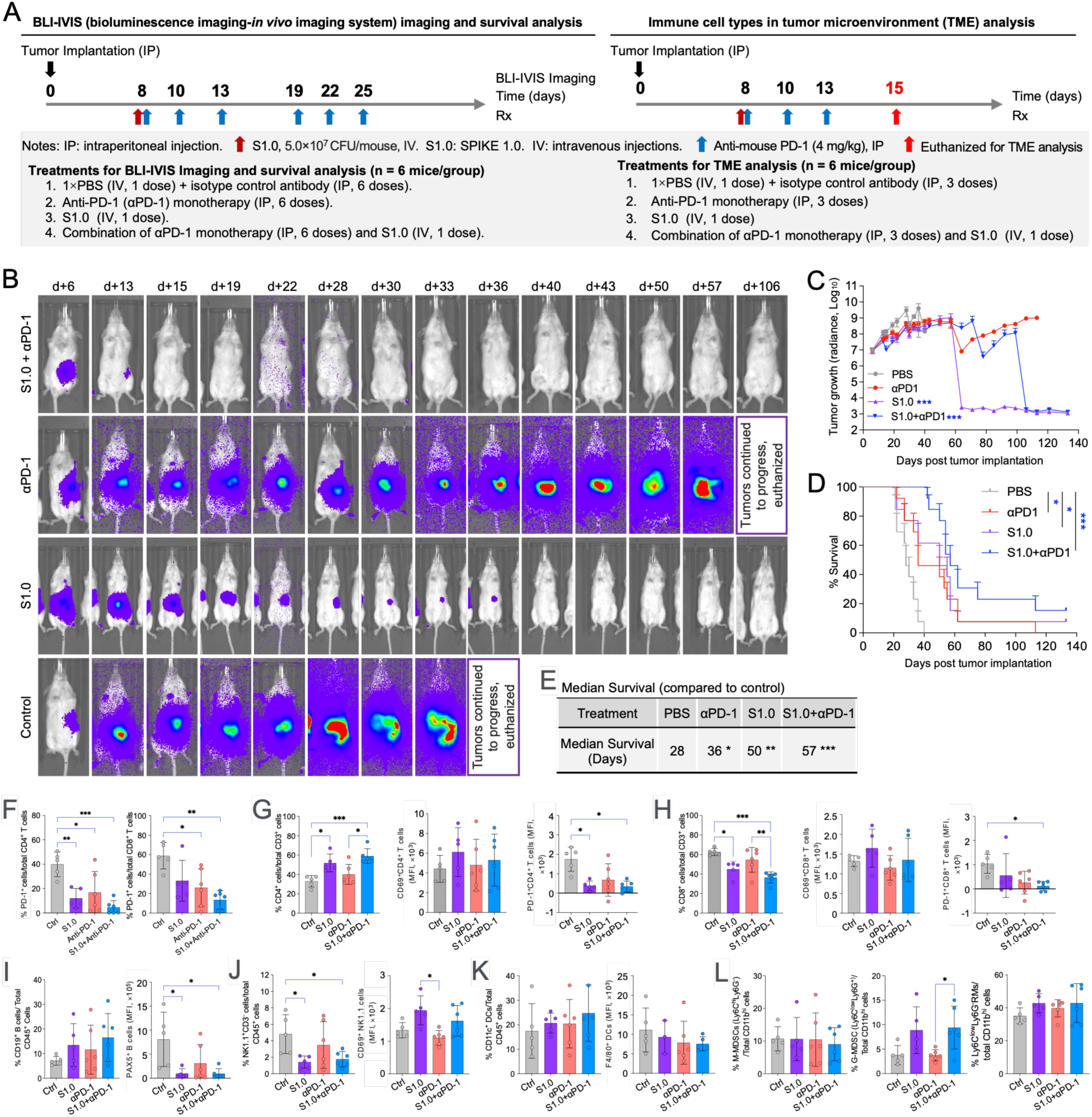
SPIKE 1.0 increases survival rate of orthotopic pancreatic ductal adenocarcinoma (PDAC) tumor-bearing mice and differentially regulates immune cell populations in TME. (A) Schematic illustration of tumor implantation, treatment schedule, treatment-efficiency evaluation, and the TME analysis in KPC-Luc orthotopic PDAC tumor mouse model. (B) Representative *in vivo* images showing monitorization of KPC-Luc PDAC tumor growth in tumor-bearing mice (n = 6 mice/group.) with the indicated treatments. d: Days post-surgery. (C) Quantification of luciferase activity showing PDAC tumor mass growth in treated mice. (D) Survival curves of the treated KPC-luciferase PDAC tumor-bearing mice. (E) S1.0 administration increases median survival of PDAC tumor-bearing mice. (F) S1.0 administration reduce % of PD-1^+^ cells to total CD4^+^ and CD8^+^ T cells in TME of PDAC mouse model. (G-L) SPIKE 1.0 differentially regulates the populations of CD4^+^ T cells (G), CD8^+^ T cells (H), terminal differentiation of B cells and long-lived plasma cells (I), NK cells (J), CD11c^+^ dendritic cells (DC) (K), and myeloid derived suppressor cells (MDSC) (L) in the TME. MFI: mean fluorescence intensity. n = 3. *, **, ***: p < 0.05, 0.01, and 0.001, respectively.

To benchmark our immunotherapeutic approach, we compared S1.0 efficacy to established immunotherapies, including chimeric antigen receipt-T (CAR-T) (mesothelin; AsPC-1), T cell receptor T (TCR-T) (NY-ESO1; H1975) cells, and ICI αPD-1 (KPC-Luc orthotopic PDAC) that target antigens expressed on the corresponding tumor surfaces (Figure 2; Figure 3A-E). We found that in these tested models, S1.0 treatment performed as well or better than corresponding CAR-T, TCR-T, or ICI benchmark immunotherapies in limiting tumor growth and extending the survival of tumor-bearing mice (Figure 2C-J; Figure 3B-E). Moreover, S1.0 worked synergistically with benchmark therapies in treating these tumors and significantly prolonged lifespan of tumor-bearing mice (Figure 2C-J; Figure 3B-E). Strikingly, complete tumor regression was achieved in a subset of mice across all tumor models treated with S1.0. These findings underscore the translational potential of S1.0 as a broadly effective microbial immunotherapy capable of inducing durable antitumor responses.

### S1.0 promotes immune cell infiltration into the TME of tumor-bearing mice

To dissect the molecular and cellular mechanisms by which S1.0 inhibited tumor growth and increased survival of tumor-bearing mice in divergent tumor models, we measured the responses of immune cells in the TME. Flow-cytometric analysis indicated that S1.0 treatment promoted reductions in PD-1 expression both independently and in combination with αPD-1 antibody in CD4^+^ T and CD8^+^ T cells isolated from the KPC-Luc orthotopic PDAC mouse model (Figure 3F). This treatment also increased immune cell infiltration into the TME (Figure 3G-L). Importantly, data from replicate experiments performed at an independent experimental site also demonstrated that in the same KPC-Luc orthotopic PDAC mouse model, SC treatment significantly promoted CD8^+^ T cell infiltration into the TME of SC-treated tumor-bearing mice (Figure 4A). These concordant findings improved the rigor of the research findings. While S1.0 treatment did not uniformly increase the frequency of tumor-infiltrating CD8⁺ T cells or NK cells, it significantly reduced PD-1 expression on CD8⁺ T cells in combination with anti-PD-1 therapy (Fig. 3F) and enhanced CD69 expression (Fig. 3H), suggesting decreased exhaustion and increased activation. These data therefore suggested that S1.0 primarily improves the functional quality of tumor-infiltrating lymphocytes rather than driving numerical expansion.

**Figure 4.**
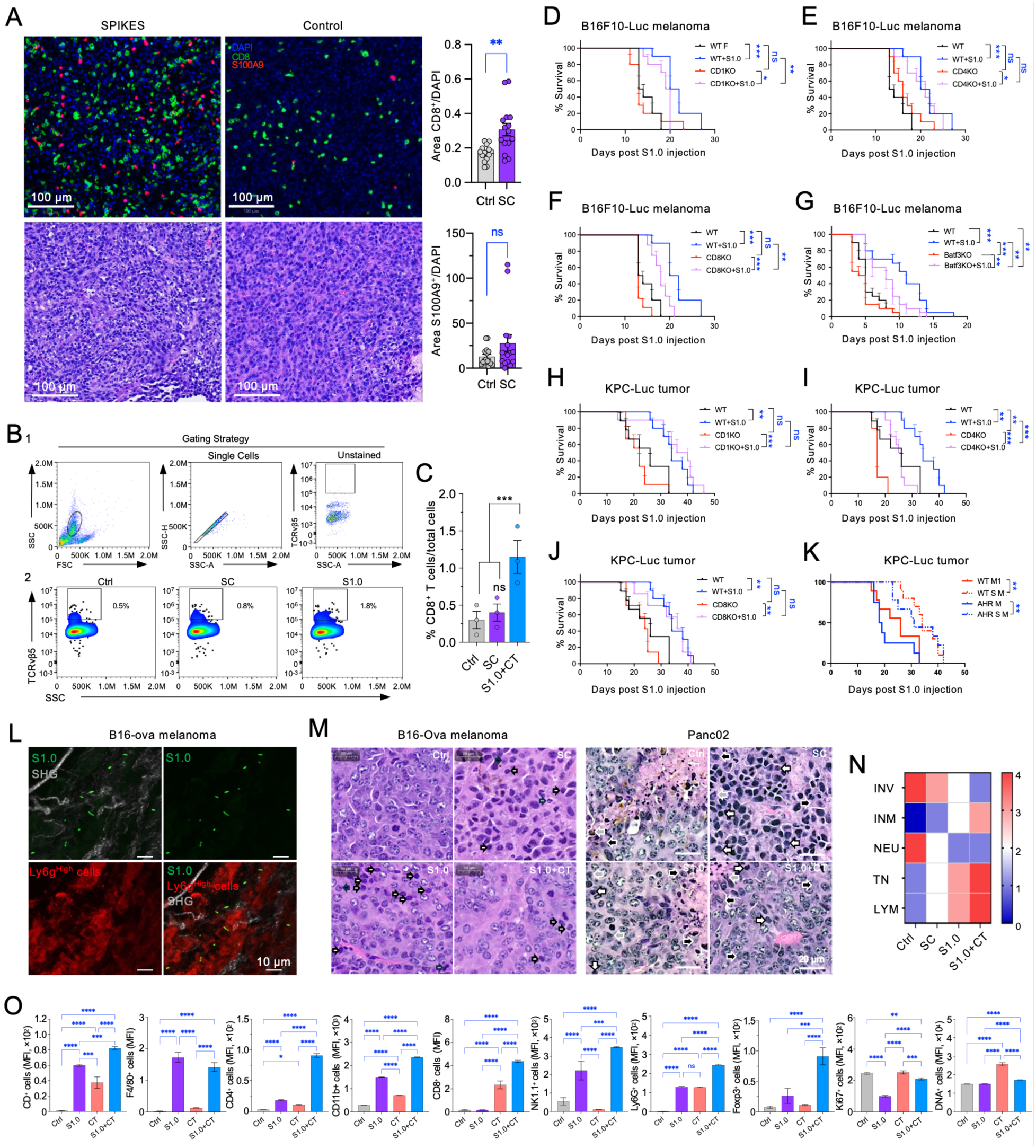
S1.0 increases immune cell infiltration into tumor microenvironments (TME) of tumor-bearing mice. (A) Confocal image analysis of CD8^+^ T cell and S100A9^+^ MDSC infiltration into orthotopic PDAC tumors in C57BL/6 mice at 3 days post S1.0 administration (n=15, 5×10^7^ cfu/mouse, upper left panel) and quantification of the indicated cell population in the TME (right panel). Lower left panel: Hematoxylin and eosin (H&E) staining analysis of explanted tumor sections from tumor-bearing mice. No difference was detected at this time point. (B) Representative gating strategy (B1) and flowcytometric analysis (B2) of OT-1 CD8^+^ T cell infiltration into B16-Ova melanoma tumors in C57BL/6 mice. (C) Quantification of flowcytometric analysis shown in (B). (D-K) CD1, CD4, CD8, Batf3, or AhR (aryl hydrocarbon receptor) play a limited role in SPIKE 1.0-mediated increase of survival of tumor-bearing mice. S1.0 increases survival of tumor-bearing CD1 (D), CD4 (E), CD8 (F), and Batf3 (G) gene knockout (KO) mice pre-inoculated with B16F10-Luc tumor cells or CD1 (H), CD4 (I), CD8 (J), and AhR macrophage (K) KO mice pre-inoculated with KPC-luciferase (KPC-Luc) tumor cells. n = 15 mice/group. (L) Multiphoton microscopic analysis of S1.0 localization in TME (green: S1.0 bacteria; red: Ly6g^+^ cells) of explanted B16-Ova melanoma (white: SHG-second harmonic generation for collagen structure). (M) H&E staining analysis of explanted tumor sections from tumor-bearing mice. White open arrows indicate lymphocytes, closed/black arrows macrophage, gray arrows neutrophil infiltration in tumor sections induced by the indicated treatments. n = 5 mice/treatment. (N) Heatmap showing the difference in the tumors with different treatments. INV: tumor local invasion, MAC: intratumor macrophage/monocyte infiltration, NEU: intratumor neutrophil infiltration, TN: tumor necrosis, LYM: intratumor lymphocyte infiltration. (O) Quantification of immune cells infiltration into the TME obtained by multiparametric spatial image analysis. Quantitative data represent mean ± SEM from three independent experiments. *, **, ***, and ****: p ≤ 0.05, 0.01, 0.001, 0.0001, respectively.

Similarly, in the B16-Ova melanoma mouse model, we found that the infiltration of OT-1 CD8^+^ T cells into the TME increased significantly (11-fold) compared to controls (Figure 4B-C), suggesting that CD8^+^ T cells may play important roles in immunomodulation and antitumor activity in the intrinsic immune dependencies of S1.0. We thus tested the antitumor roles of CD8^+^ T cells and other immune cell types in the presence of S1.0 using gene knockout (KO) mice that are deficient in functional NK cells (CD1 KO), CD4^+^ T cells (CD4 KO), CD8^+^ T cells (CD8 KO), CD8α^+^ dendritic cells (DCs) (Batf3 KO), which are crucial for priming CD8^+^ T cells and initiating effective immune responses against tumors^24,25^, and AhR (aryl hydrocarbon receptor) macrophages (AhR-KO macrophages). We treated wild-type (WT) or gene KO mice carrying B16F1-Luc syngeneic murine melanoma or KPC-Luc orthotopic PDAC tumors with S1.0. We found that treatment generally led to significantly prolonged survival compared to mock-treated controls (Figure 4D-K) in these immune-deficient models. Notably, treatment of CD8 KO mice with either B16F10 (Figure 4F) or KPC-Luc (Figure 4J) tumors, or similar treatment of CD4 KO mice with KPC-Luc (Figure 4I) resulted in reduced survival of tumor-bearing mice compared to their respective WT controls. These findings suggest that individual immune cell populations (e.g., CD4⁺ or CD8⁺ T cells) contribute to the magnitude of the therapeutic response by S1.0. These findings also supported the hypothesis that S1.0 engages multiple immune functions in context-dependent fashion, rather than relying upon a single immune subset. Collectively, our findings indicated that the depletion of the functions of any one of the interrogated immune cell types did not thwart the anticancer effect of S1.0.

Multi-photon microscopy analysis of subcutaneously implanted B16-Ova tumors showed that S1.0 localized in a region associated with high-expression level of Ly6g^+^ cells (Figure 4L). Our hematoxylin and eosin (H&E) staining analysis showed increased intraneoplasm macrophages, lymphocyte infiltration, and tumor necrosis, but reduced tumor local invasion in mice treated with S1.0 or a combination of S1.0 and antigen specific CAR-T cells (Figure 4M-N). Importantly, these findings were validated by the multiparametric spatial imaging analysis, which showed increased infiltration of F4/80^+^ macrophages, CD8^+^ T cells, CD4^+^ T cells, NK 1.1 cells, and other immune cells into the TME of B16-Ova melanoma tumors (Figure 4O, Figure S3). Collectively, our findings demonstrated that S1.0 treatment limited tumor growth and even led to regression in tumor size in some mice. In all the tumor models tested, S1.0 treatment enhanced and prolonged survival of tumor-bearing mice, consistent with the observed increases in immune cell infiltration into the TME of tumor-bearing mice.

### S1.0 promotes inflammatory activity and CD8^+^ T cell infiltration in humanized mice

To assess the translational relevance of S1.0, we performed studies using humanized mouse models of lung cancer (H1975) and Merkel cell carcinoma (MCC13) (Figure 5A). In these models, the NSG mice were first engrafted with human CD34^+^ hematopoietic progenitor cells, resulting in animals that carried human B cell, T cell, CD11c+ dendritic cell and other monocyte populations^26–28^. Our findings demonstrated that S1.0 intervention substantially reduced tumor growth and increased the survival of the treated animals (Figure 5B-E). Moreover, S1.0 treatment alone and with TCR-T cells also resulted in minimal weight loss in tumor-bearing mice compared to untreated, SC treated or TCR-T alone treated mice, suggesting that the S1.0 intervention, alone or in combination with TCR-T cells was well-tolerated and significantly improved the outcomes (Figure 5F). These pre-clinical data therefore supported the human-relevance of the observations in lung and skin cancer. Consistent with findings from syngeneic mouse model studies, we observed significantly increased infiltration of immune cells (Figure 5G-I, K), along with reduced expression of PD-1 (Figure 5J, K) in the TME of humanized mice treated with S1.0 and TCR-T (Figure 5G-K). Similarly, H&E staining analysis demonstrated that S1.0 treatment increased infiltration of immune cells, including macrophages and lymphocytes, and tumor necrosis in S1.0 or combination treatments (Figure 5L-M). Collectively, these results demonstrated that in the humanized mouse model, S1.0 significantly promoted infiltration of immune cells, reduced tumor growth, and improved survival of tumor-bearing mice, indicating the versatility of our engineered bacteria.

**Figure 5.**
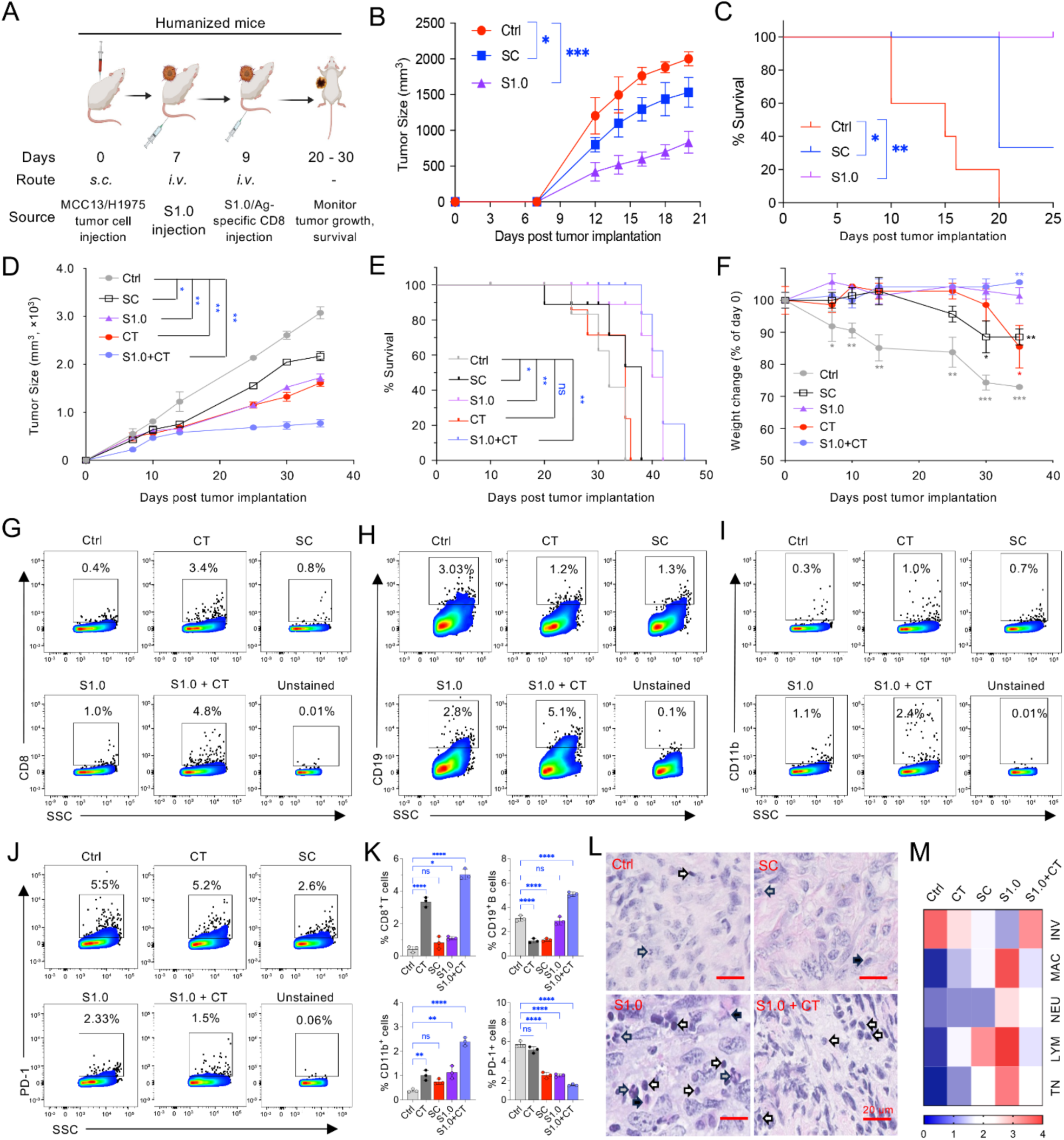
S1.0 promotes inflammatory immune activity and CD8^+^ T cell infiltration in humanized mice. (A) Schematic experimental design of humanized-mouse MCC13/H1975 tumor model. (B) MCC13 tumor growth curves. (C) Survival rate of MCC13 tumor-bearing mice. (D) H1975 tumor growth curves in tumor-bearing mice. (E-F) Survival curves (E) and relative weight change (F) of H1975 tumor-bearing mice. (G-I), CD8^+^ T cells (G), CD19^+^ B cells (H), and CD11b^+^ monocytes (I) infiltration into tumors shown by flowcytometric analysis. (J) PD-1^+^ cells in the TME shown by flowcytometric analysis. (K) Quantification of flowcytometric analysis showing in panels (G) to (J). (L) H&E image analysis of explanted tumors. Representative images are shown. Open white arrows: lymphocytes, closed black arrows: macrophages, gray arrows: neutrophils. Bars: 20 μm. (M) Heatmap analysis of H&E scoring in tumor sections showing as a representative in panel (L). INV: local invasion, MAC: macrophage, NEU: neutrophils, TN: tumor necrosis, LYM: lymphocytes. Quantitative data represents mean ± SEM from three independent experiments. *, **, and ***: *p* < 0.05, 0.01, and 0.001, respectively.

### S1.0 upregulates chemokine signaling and the expression of AA receptors on T cells

The B16-Ova melanoma tumor-mouse model is considered a useful tool in cancer immunotherapy research because it allows for the precise tracking of immune responses against a defined ovalbumin-antigen in an aggressive, poorly immunogenic B16F10 cell line. The model has been widely deployed in immunocompetent C57BL/6 mice to study ICIs, cancer vaccines, and ACT^29,30^, and therefore was useful for our studies. To elucidate the molecular mechanisms by which S1.0 inhibited tumor progression and remodeled the cellular immune profile of the TME, we performed single-cell RNA sequencing (scRNA-seq) analyses of explanted murine B16-Ova melanoma tumors on day 21 and day 28-post tumor cell implantation (PTCI) (Figure S4A). Using cell type-specific canonical gene markers (Figure S4B-E; Table S1) and the Seurat pipeline^31^, we performed clustering analysis to identify the abundance of assorted cell types in the TME (Figure 6A; Figure S4F-I). scRNA-seq analysis identified hundreds of differentially expressed genes (DEGs) in each immune cell type in tumors treated with S1.0 compared to controls (Table S2, S3). Further analysis of these DEGs (Figures 6; S5; S6; Table S2, S3) showed that S1.0 significantly increased infiltration of many immune cells into the TME (Figure 6B). At day 21 PTCI, we also observed differential expression of genes encoding antigen presentation proteins such as H2Aa1 and H2Ab1 (Figure 6C-D) and chemokine ligands like Ccl3 and Ccl4 (Figure 6E) in macrophages of S1.0 treated tumors compared to controls. Moreover, we found that the gene expression of chemokine and other immune receptors also increased in S1.0-treated tumors. For example, CD69, a biomarker that not only is rapidly expressed upon immune activation but also acts as a co-stimulatory molecule for T-cell activation, proliferation, and natural killer (NK) cell cytotoxicity^32^ was upregulated in S1.0 treated mice (Figure 6F). Interestingly, S1.0 also promoted Ccr7 expression, a marker for central memory T cells (T_cm_), whereas S1.0 in combination with CAR-T significantly promoted effector memory T cells (T_em_) by increasing the expression of the Ccr5 chemokine receptor marker (Figure 6F). While the data suggested that S1.0 increased macrophage activation T_cm_ and T_em_, recruitment to the TME in the B16-Ova model, the extent to which these effects drove the enhanced chemotaxis of adaptive immune cells to the TME in other tumor models requires further clarification.

**Figure 6.**
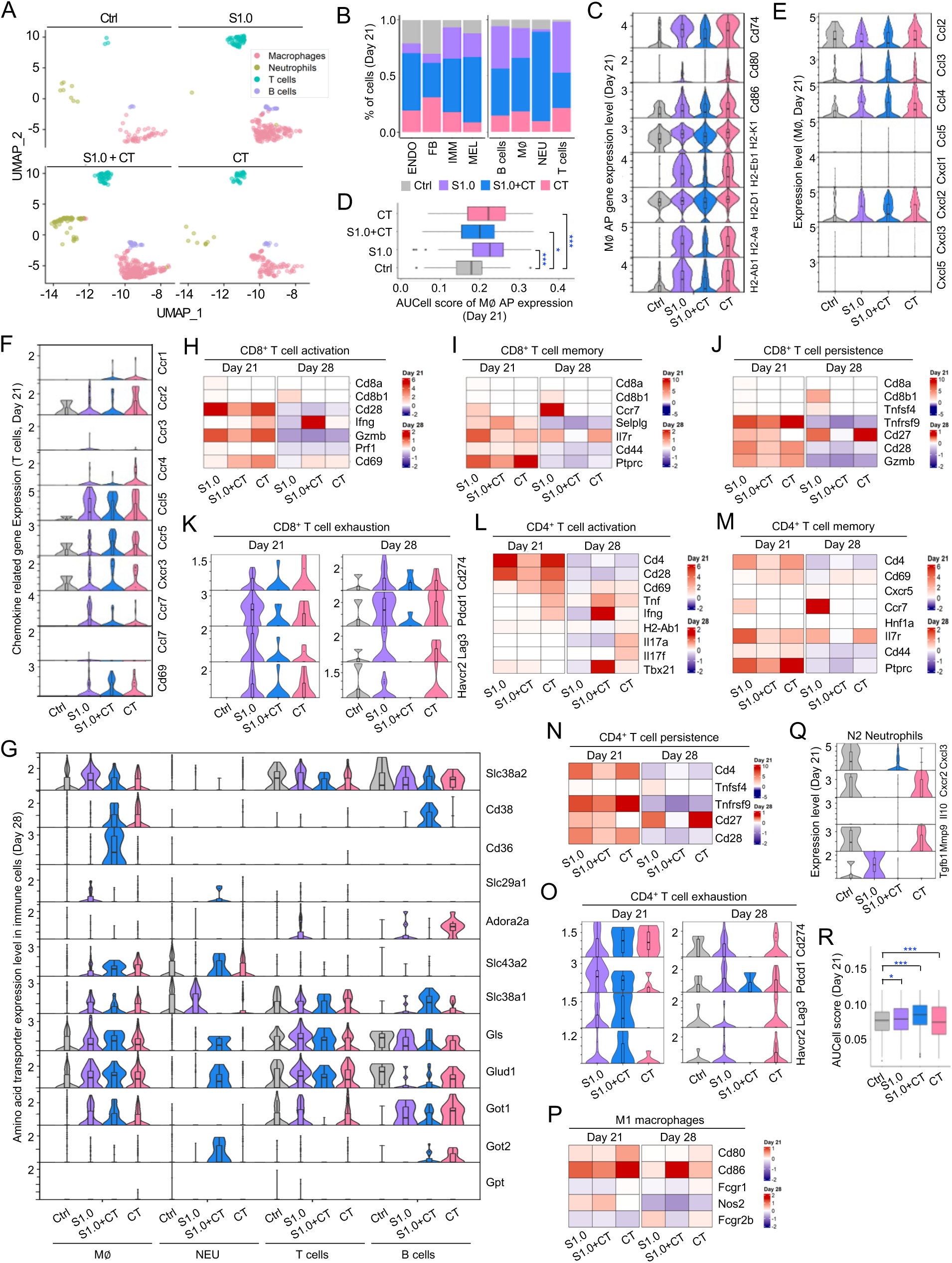
S1.0 upregulates biomarkers of immune cell activation and modulates amino acid (AA) receptors in immune cells in TME of B16-Ova melanoma tumor-bearing mice. (A) Representative UMAP (uniform manifold approximation and projection) clustering analysis of the indicated immune cells in the TME at day 21 PTCI of tumor-bearing mice with the indicated treatments. (B) Quantification of all cells and specific immune cells in the TME. (C) Representative violin plots showing the expression of antigen presentation genes and co-stimulatory markers on macrophages (M∅∥). (D) AUCell score analysis of antigen presentation and co-stimulation of macrophages shown in (C). (E-F) Representative plots of chemokine ligands expression on macrophages (E) and chemokine receptor expression on T cells (F). (G) AA receptors and transporter marker gene expression on the indicated immune cells. (H-J) Heatmap showing expression of marker genes indicating CD8^+^ T cell activation (H), CD8^+^ T cell memory (I), and CD8^+^ T cell persistence (J). (K) Violin plots showing the expression of CD8^+^ T cell exhaustion marker genes. (L-M) Heatmap analyses of expression of markers showing CD4^+^ T cell activation (L), CD4^+^ T cell memory (M), and CD4^+^ T cell persistence (N). (O) Violin plots showing the expression of CD4^+^ T cell exhaustion marker gene expression. (P) Heatmap showing expression of markers identifying M1 macrophages. (Q) violin plots showing the expression of marker genes that identify N2 neutrophil in TME at day 21 PTCI. n = 3. (R) AUCell score analysis of apoptosis-associated genes in melanocytes on day 21 post tumor-cell implantation (PTCI). n=3. *, ***: signification at p < 0.05 and 0.001, respectively.

AAs are essential nutrients for the survival and growth of immune and cancer cells. Interestingly, scRNA-seq and AA metabolism enrichment analyses in different cell types at day 21 PTCI indicated that in the TME of S1.0-treated tumors, global AA metabolism, including alanine and aspartate metabolism, was selectively upregulated in T cells (Figure S5; Table S2, S3). The expression of AA transporters was also significantly increased in T cells (Figure 6G). Importantly, AA transporters Slc38a1 and Slc38a2, two members of the solute carrier family 38 (SLC38) involved in the transport of glutamine and other AAs such as alanine, serine, proline, and glutamine into cells^33,34^, along with glutamate dehydrogenase 1 (Glud1) and glutamate oxaloacetate transaminase 1 (Got1), were significantly upregulated (Figure 6G). The upregulation of these proteins was sustained in tumor-bearing mice treated with S1.0 and CAR-T cells (Figure 6G). These findings suggested that S1.0 may metabolically reprogram T cells in the TME by the functional modulation of AA-associated pathways, potentially mitigating early exhaustion and enhancing effector functions. These findings underscored the cell-type specificity of S1.0-induced metabolic reprogramming.

To investigate whether enhanced AA utilization and metabolic rewiring in CD4^+^ and CD8^+^ T cells translated into increased functional capacity within the TME, we analyzed their activation, persistence, memory formation, and suppressive capabilities. Our scRNA-seq analysis (Figure 6H-R; Table S2, S3) revealed a marked increase in CD8^+^ T cell activation in the S1.0-treated tumors at day 21 PTCI (Figure 6H). Notably, S1.0-treated CD8^+^ T cells demonstrated an enhanced effector/central memory phenotype (Figure 6I), which was attributed to reduced activation-induced cell death and a prolonged persistence of functional CD8^+^ T cells (Figure 6J). In contrast, the initial heightened activation observed in the antigen-specific CAR-T-treated group appeared to contribute to early exhaustion of CD8^+^ T cells (Figure 6J-K). Importantly, S1.0 alone, and the combination of S1.0 and CAR-T treatments, induced a distinct population of tumor-associated CD8^+^ T cells characterized by a Ccr5^high^Il7^high^Gzmb^high^ Cd27^high^Cd28^high^ phenotype on day 21 PTCI (Figure 6F, H-J), suggesting robust and sustained anti-tumor activity. We observed similar phenotypes in tumor-associated CD4^+^ T cells with the gene expression profile indicating lower but sustained initial activation, higher memory phenotypes, higher persistence and lower suppressive capacity on day 28 PTCI, compared to controls (Figure 6L-O). Consistent with the hypothesis, we observed reduced expression of exhaustion markers PD-1 (Cd274), Tim-3 (Havcr2) and Lag3 and increased IL-7 signaling in both CD4^+^ and CD8^+^ T cells in the combination S1.0 and CAR-T treatment compared to the S1.0 or CAR-T treatment alone (Figure 6I, K, M, O). These findings suggested their enhanced functional states, particularly on day 28 PTCI, with significant differences observed in Tim-3 and Lag-3 expression (Figure 6I, K, M, O). Furthermore, we observed upregulated proinflammatory M1 macrophages (Figure 6P), B cell activity (Figure S6A-C), and AhR signaling in macrophages and T cells (Figure S6D-E) in tumors of S1.0, or combination of S1.0 and CAR-T, treated mice. The downregulation of cell markers associated with tumor promotion, including those of M2 macrophages, myeloid-derived suppressor cells (MDSCs), and N2 neutrophils, all of which are pro-tumorigenic, strongly immunosuppressive, and promote tumor angiogenesis and metastasis^35–37^, was also observed in the tumors of S1.0-treated mice (Figure 6Q; Figure S6F-I). Finally, we observed upregulated apoptosis-associated genes (Figure 6R), implying more apoptosis in melanocytes in the TME of S1.0-treated tumors, consistent with the downregulation of marker genes associated with tumor-promotion.

Overall, our scRNA-seq analysis suggested that S1.0 promoted the infiltration of immune cells into the TME and increased antigen presentation and chemokine expression activity in innate immune cells, thereby resulting in the increased functional capacities of CD8^+^ and CD4^+^ T cells. Furthermore, S1.0 treatment selectively promoted CD8^+^ and CD4^+^ T cell increases in the expression of genes that control AA metabolism.

### S1.0 remodels AA metabolic pathways

The regulation of AA receptors and transporters in CD8^+^ and CD4^+^ T cells by S1.0 encouraged us to perform a metabolomics analysis of the TME in S1.0– and control-treated tumor-bearing mice. The partial least squares-discriminant analysis (PLS-DA) score plots based on the TME metabolome database revealed significant separation of all the treatments (Figure 7A). Metabolite analysis demonstrated increases in phosphatidylcholine, phosphatidylserine, phosphatidylethanolamine, and methylthioadenosine, a naturally occurring sulfur-containing nucleoside that suppresses tumors by inhibiting tumor cell proliferation, invasion, and the induction of apoptosis^38,39^ (Figure 7B; Table S4). Importantly, S1.0 treatment significantly increased the levels of the mixed 3-, 4-, 5-, 6-, and 7-HIs in the TME compared to the control group, where these metabolites fell below the detection limit (Figure 7C; Table S4). We observed that metabolites associated with tumor proliferation and growth, including lactate, glutamine and glutamate^40^ significantly decreased in the TME of tumors from S1.0-treated mice (Figure 7D; Table S4). Metabolic Set Enrichment Analysis (MSEA) revealed that in the S1.0-treated TME, metabolic pathways associated with tumor cell development, proliferation and metastasis, including arginine biosynthesis^41^, glycolysis or gluconeogenesis^42^, alanine, aspartate and glutamate metabolism^43^, were dramatically modulated (Figure 7E-F, Figure S7A; Table S2-S4). Moreover, scRNA-seq gene set enrichment analysis demonstrated the upregulation of genes involved in glycolysis and down-regulation of genes in arginine metabolism pathways in CD4^+^ T cells (Figure S7B-C) on day 21 and day 28 PTCI, respectively. These transcriptomic results further supported findings from the metabolomics studies.

**Figure 7.**
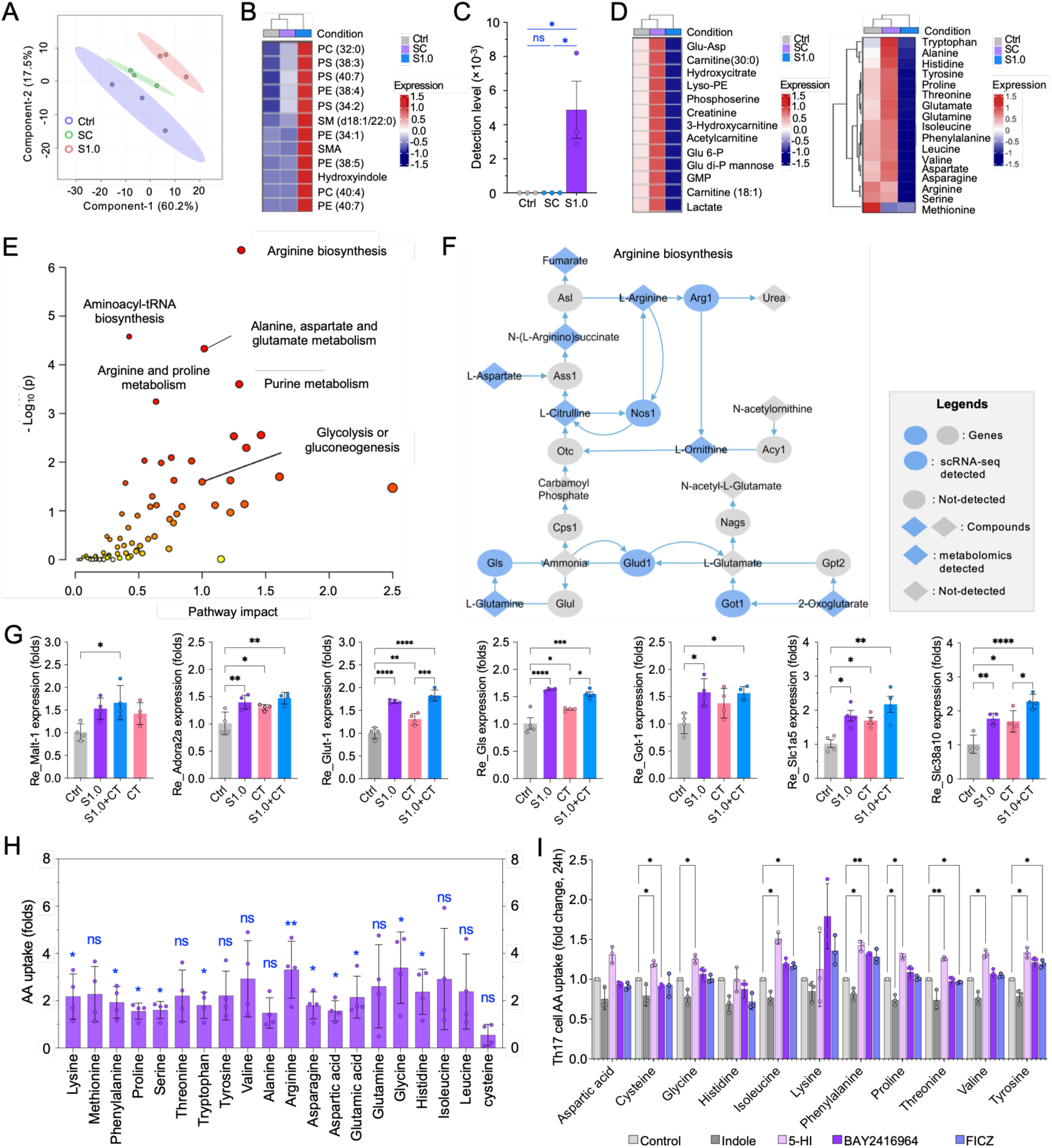
S1.0 remodulates AA metabolic pathways impacting tumor metabolome. (A) Partial least squares-discriminant analysis (PLS-DA) of the metabolome of control, SC and S1.0 treated B16-Ova tumors. (B) Comparison of metabolite levels of phospholipids-PC (phosphatidylcholine), PS (phosphatidylserine), PE (phosphatidylethanolamine), HI, and metabolite intermediates of glycolysis from the TME at day 21 PTCI of the treated tumors. (C) Detection levels of the mixed 3– to 7-HI in the TME of S1.0– and control-treated tumor bearing mice. (D) Representative heatmap of the comparative metabolomics analysis of amino acids. (E) Representative pathway impact analysis derived from the metabolomics profile. (F) Network map analysis of arginine biosynthesis pathway in the TEM of B16-Ova tumor treated with Ctrl, SC or S1.0 at Day 21. The analyzed datasets are derived from the combined scRNA-seq data (Table S2, S3) and metabolomics (Table S4). (G) qPCR validation of the upregulation of AA receptors and transporter markers in immune cells in TME of B16-Ova melanoma tumor-bearing mice treated with the indicated treatments at 21 days PTCI. (H) AA uptake profiling of T cells in TME of B16-Ova melanoma tumors treated with control or SC. AA uptake data of control was normalized as 1. (I) Th17 cell AA uptake profiling analysis at 24 h post co-culture with 5-HI and other small molecules as indicated. *: *p* < 0.05, **: *p* < 0.01, ***: *p* < 0.001, ****: *p* < 0.0001.

To validate findings from the scRNA-seq and metabolomics analyses, we performed quantitative real-time PCR (qPCR) and *ex vivo* AA uptake profiling assays to analyze the expression of some key identified AA transporters and the AA uptake levels of T cells in the TME from tumor-bearing mice treated with S1.0 or control. As expected, we found that the expression of the tested AA transporters was significantly upregulated (Figure 7G) and that the uptake of the tested AAs selectively increased (Figure 7H) in the TME of S1.0– or SC-treated B16-Ova melanoma tumors.

To determine if 5-HI and AhR signaling affect immune cell AA uptake, we cocultured T cells isolated from mouse spleens or Th17 cells (differentiated from the T cells) with 5-HI and other small molecules, include indole, BAY 2416964 (AhR antagonist)^44^, and FICZ (AhR agonist)^45^, and determined AA uptake by the small molecule-treated cells at 24 hours post treatment (hpt). We found that 5-HI significantly promoted T cell AA uptake at 24 hpt (Figure S7D). Importantly, after T cells differentiation into Th17 cells, 5-HI also facilitated AA uptake by the differentiated Th17 cells (Figure 7I). These findings demonstrate that 5-HI contributed, at least partially, the increased AA uptake by immune cells, and imply that the metabolically engineered strain S1.0 that constitutively produces 5-HI enhances immune cells activation and maintains their functional immune activity activates in the TME via increasing their AA uptake.

Collectively, our metabolomics and scRNA-seq analyses demonstrated that metabolites and/or metabolic synthesis pathways related to immunomodulation increased and those associated with tumor cell growth and metastasis decreased in the S1.0-treated TME, thereby revealing metabolic rewiring of immune cells in the TME.

## Discussion

This study shows that a single systemic dose of the engineered bacterial therapeutic S1.0 induces strong antitumor responses in multiple murine and humanized cancer models by reprogramming the immunosuppressive TME. S1.0 created a pro-inflammatory, tumoricidal environment while maintaining a favorable safety profile. It activated both innate and adaptive immune responses, as indicated by increased levels of TNF-α, perforin, and GrB. S1.0 overcomes immune exclusion in cold tumors and restored function in exhausted T cells within hot tumors. These results establish S1.0 as a potent microbial immunotherapy candidate that broadly enhances antitumor immunity by remodeling immune activity within solid tumors. The ability of 5-HI to upregulate CD38 expression in macrophages and enhance immune cell AA uptake further underscores its role in reshaping tumor metabolism, a novel strategy that directly links bacterial metabolic engineering with immune potentiation. Notably, S1.0 outperformed its parental strain in tumor cell cytotoxicity without compromising bacterial invasion efficiency, highlighting the precision of its engineered immunostimulatory functions. Moreover, the observed enhancement of antigen-specific human CD8^+^ T cell cytotoxicity against lung cancer cells suggests the translational potential of this platform. These findings position S1.0 as a potential bacterial immunotherapy with the capacity to remodel immune landscapes across diverse tumor types, setting the stage for innovative metabolic-immune interventions in oncology.

S1.0 carries *Tna* and *Tmo* that together can produce metabolites such as HIs, including 5-HI. In this work, we focused on the analysis of HIs (5-HI was used as an example), which were consistently detected and functionally associated with the observed immunomodulatory effects. However, Tna can contribute to the synthesis of additional indole/tryptophan-derived metabolites (e.g., I3A, IAA, IPA) with multiple immune-modulating functions; Tmo-associated pathways may yield p-cresol and related aromatic metabolites. Therefore, we cannot rule out the immunomodulation contributions of additional metabolic products by S1.0. The bioactive features of other metabolites produced by S1.0 require further investigations.

Besides the use of the well-characterized and responsive B16-Ova melanoma model to define candidate molecules and pathways engaged by S1.0 treatment, we extended our studies to human solid tumor xenograft models using lung cancer (H1975) and pancreatic cancer (AsPC-1) cells implanted heterotopically in immunocompromised NSG mice to assess the relevance of S1.0 beyond murine syngeneic settings. The NSG model was particularly well-suited for this purpose: the absence of endogenous B, T, and NK cells minimized confounding host adaptive immune responses as well as tolerance of human tissue explants. This model therefore enabled accurate evaluation of how S1.0 modulated the TME of human-derived tumors in a living system. S1.0 treatment significantly reduced tumor burden and extended survival across both syngeneic (B16-OVA melanoma, Panc02) and xenograft (H1975, AsPC-1) models as well as humanized mouse models of lung cancer (H1975) and Merkel cell carcinoma (MCC13). S1.0 outperformed traditional checkpoint inhibitors, synergized with chemotherapy, and worked synergistically with the benchmark therapies in treating the tested tumors to significantly prolong survival of tumor-bearing mice. Complete tumor regression was achieved in a subset of mice across all tumor models treated with S1.0. Findings from xenograft mouse models (NSG background) also suggested that S1.0 may exert partial effects in the absence of adaptive immunity, potentially mediated through residual host components such as myeloid-lineage cells or direct bacterial modulation of the TME.

S1.0 reshaped the suppressive TME through broad immunomodulatory effects rather than reliance on a single immune cell type. Across tumor models, we observed notable differences in immune cell composition and infiltration, reflecting intrinsic tumor-specific constraints on immune accessibility and function. Despite this heterogeneity, S1.0 consistently promoted metabolic reprogramming and functional activation of immune populations within the TME. scRNA-seq analysis revealed increased infiltration of innate and adaptive immune cells, including CD8⁺ and CD4⁺ T cells, central and effector memory T cells, macrophages, and N1 neutrophils, although the relative contributions of these populations varied by tumor type. The efficacy of S1.0 has been demonstrated across all our tested tumor models. However, findings from a single tumor model are not universally generalizable and require further validation in additional contexts. Therefore, the detailed mechanisms by which S1.0 modulates the TME in more immunosuppressive and translationally relevant models require further investigation.

AhR is a key immunological regulator in the TME. AhR signaling is widely recognized as a “double-edged sword” or “context-dependent regulator” that does not uniformly promote antitumor responses, but rather drives both immunosuppressive and, in some settings, inflammatory phenotypes in T cells and myeloid cells depending on the specific ligand and TME cues^46–48^. Given the role of the AhR as a central regulator of immune responses, our data support a model in which indole derivatives such as 5-HI drive AhR-dependent immune reprogramming across multiple compartments, leading to enhanced cytotoxic programs, memory associated features, and recruitment of effector CXCR3⁺ CD8⁺ T cells. These effects were reinforced by increased antigen presentation and chemokine expression in macrophages.

While regulatory T cells increased in certain contexts, this likely reflects compensatory regulation in response to heightened immune activation rather than dominant immunosuppression, as antitumor efficacy was maintained. For example, in the B16-OVA tumor model, an increased Treg population was observed in the S1.0+CT treatment (Figure 4O). Treg expansion is a well-recognized consequence of heightened immune activation and inflammation within the TME, where regulatory mechanisms are engaged to limit excessive tissue damage^49^. Importantly, despite the increased Treg population, S1.0+CT treatment resulted in overall improvement of the antitumor landscape, including concurrent enhancement of effector T cell activity and inflammatory signaling. Therefore, Treg expansion likely reflects a compensatory regulatory response rather than dominant immunosuppression. Consistent with this systems-level rewiring of immune functions, S1.0 reduced pro-tumorigenic N2 neutrophils (Fig. 6Q) and induced only modest tumor control in immunodeficient NSG mice, supporting the conclusion that S1.0 functions primarily by remodeling the TME to potentiate adaptive cellular therapies rather than through immune cell–intrinsic or immune-independent mechanisms.

The metabolic reprogramming induced by S1.0 may represent another significant mechanism by which this engineered strain enhances anti-cancer immunity. scRNA-seq and metabolomics findings demonstrate that the expression and activity of AA receptors and transporters increased in the TEM of tumors treated with S1.0. Our data support the hypothesis that T cells from the TME of tumor-bearing mice treated with S1.0 increase their AA uptake. AA availability, critical for immune cell function^50,51^, was markedly altered in the TME of S1.0-treated mice. Key transporters such as Slc38a1 and Slc38a2, were upregulated in CD4^+^ and CD8^+^ T cells, supporting their activation, proliferation, and effector functions under nutrient-deprived conditions^33,34^. These observations align with previous reports linking AA metabolism (e.g., glutamine, branched-chain amino acids, methionine) to enhanced T cell activation, proliferation, and functionality^52–56^. Elevated expression of Glud1 and Got1 (Figure 6G), which facilitates glutaminolysis^57^ and aspartate metabolism respectively, underscores the metabolic adaptations of immune cells to the hostile TME^58^. These metabolic pathways and metabolites may not only sustain cytokine production and cytotoxic activity but also promote the formation of memory T cells, which rely on mitochondrial health and oxidative phosphorylation (OXPHOS) for persistence. However, the direct link between AA metabolism and enhanced T-cell functionality requires further validation in S1.0-treated tumor models.

Additionally, the upregulation of Ccr7 and Il7r in T cells within the TME highlights their role in enhancing T cell trafficking, survival, and memory differentiation^59^. Ccr7 directs T cells to tumor-draining lymph nodes (TDLNs) for priming and expansion, while Il7r signaling supports effector and memory T cell survival in the immunosuppressive TME. These mechanisms collectively enhance the endurance and readiness of T cells to mount effective antitumor responses. Furthermore, the observed increase in Ccr5 expression and AA arginine suggests a higher potential for differentiation into effector memory phenotypes, critical for sustained immune surveillance.

5-HI exerts direct functional effects on immune cells, including enhanced cytokine production, CD8⁺ T cell cytotoxic activity, modulation of CD4⁺ T cell differentiation, and promotion of M1-associated macrophage activation (Figure 1A-F, L-M; Figure S1A, E-L). Moreover, 5-HI promotes T and Th17 cell AA uptake (Figure 7I; Figure S7D). It is noted that differences in immune cell types and/or populations exist between tumor models, resulting from a combination of intrinsic tumor differences and potential variability in metabolite dynamics. However, we cannot rule out the possibility that these differences also stem from varying HI production by S1.0 across the models since high levels of the mixed 3– to 7-HI were detected in the TME from S1.0-treated tumor bearing mice (Figure 7C). The dynamic association of the HI level (in the presence of S1.0) and AA metabolites in the TME, tumor therapeutic efficacy of HI supplementation in vivo, as well as the precise mechanisms by which HIs and S1.0 functionally immunomodulate immune cells to improve the immune-TME via metabolic reprogramming of AA need further investigation.

Collectively, our findings support a model whereby S1.0-induced immunoregulatory mechanisms contribute to cancer control. Upon administration, S1.0 homes to and then resides in the TME, which may be mediated by Ly6g^High^ MDSCs^60^. Next, S1.0 1) releases HIs, promotes antitumor chemokine production, and increases immune cell (e.g., CD8^+^ T cell) infiltration into the TME, CD4^+^ T cell polarization, and tumor cell killing; 2) activates immune cell functions, including CD8^+^ cytotoxic T lymphocyte (CTL)-directed tumor apoptosis, antigen uptake and presentation, functional M1 macrophage polarization, N1 neutrophil polarization, and CTL activation and persistence; and 3) reprograms immune cell metabolomes and increases T cell AA uptake, resulting in increased CD8^+^ T cell activation, the numbers of memory CD8^+^ T cells and their persistence and CTL activity. Finally, these immunomodulatory activities stimulated by S1.0 substantially suppress tumor growth and enhance survival. In comparison to current cellular therapies like CAR-T or TCR-T, which are often prohibitively expensive and complex, bacterial therapeutics like S1.0 may therefore offer an affordable, scalable, and efficacious alternative.

## Resource availability

### Lead contact

Further information and requests for resources and reagents should be directed to and will be fulfilled by Dr. Paul de Figueiredo.

### Materials availability

This study did not generate new unique reagents.

### Data and code availability

All data supporting the findings of this study are available in the main text and its supplementary information. Further information is available from the corresponding authors on reasonable request.

## Supporting information

Supplemental Table 1-5, and will be used for the link to file on the preprint sites

## Acknowledgments

We thank the IMIL microscopy core at Texas A&M University, College of Medicine, Steve Fulwood and Kalli Landua from Nikon, William J. Brown (Cornell University) for insightful discussions in the early phases of this work, and Hsing Fann (Texas A&M University) for the technical assistance. The study was supported by the NIH Grants No. R01CA221867, R21AI167793 and R01CA273002 to JS, the ARPA-H (No. 1AY1AX000010-01) to PDF, JS, AH, the R01CA273002 and the Roy Blunt NextGen Precision Health Endowment at the University of Missouri to PDF; the Department of Biotechnology (Grant BT/RLF/Re-entry/17/2022) and Anusandhan National Research Foundation, Govt of India (Grant ANRF/ECRG/2024/003569/LS) to JD. the Japan Society for the Promotion of Science KAKEN (23K24145, 23K18207, 22KK0112), the Japan Agency for Medical Research and Development (223fa627005,25ama221337h0002) to KSK. The content of the presented information does not necessarily reflect the position or the policy of the Government, and no official endorsement should be inferred.

## Author Contributions

Conception and design: P.dF., J.S. J.K.D. Development of the methodology: P.dF., J.S., J.K.D., S.S., Q.Q.M., C.J.T., F.G., A.C., S.S.N., C.A., M.M.K-M., S.P.C. Funding acquisition: P.dF., J.S., J.K.D., K.S.K. Experimental data collection: J.K.D., S.S., C.J.T., F.G., A.C., S.S.N., B. F., C.A., Y.A., M.M.K-M., S.P.C., A.M.R.C., K.D., A.K., E.R. Formal analyses: P.dF., J.S. J.K.D., Q.Q.M., L.G.A., S.S., A.C., S.S.N., C.J.T., S.G., J.J.C., N.N., X. Q.,S.H., A.H., K.S.K., A.J., T.A.F., R.C.A., C.Y. Writing the original draft: P.dF., J.K.D., Q.Q.M., J.S. Writing, reviewing and/or revising the manuscript: J.K.D., P.dF., J.S. QMQ, L.G.A. Study supervision: P.dF., J.S. All authors read and approved the final manuscript.

## STAR ★ METHODS

Detailed methods are provided in the online version of this paper and include the following:

- KEY RESOURCES TABLE
- METHOD DETAILS
  ○ Bacterial culture and metabolic engineering
  ○ Cell culture
  ○ Flowcytometric assessment of 5-HI activated CD8^+^ T cells and CD4^+^ T cells
  ○ *In-vitro* cytotoxic assessment of CD8^+^ T cells activity against various cancer cells
  ○ Identification of HIs from S1.0 bacterial culture filtrate
  ○ Mouse complete blood count (CBC) assays:
  ○ S1.0 biodistribution in tumor-bearing mice:
  ○ Goat S1.0 safety assessment:
  ○ CAR-T and TCR-T cell preparation
  ○ Murine tumor models
  ○ Fluorescent cytometric analysis of biomarker expression in PDAC tumor-bearing mice
  ○ Spatial multiplex immunohistochemical imaging analysis
  ○ Multiphoton image analysis
  ○ Metabolomics analysis
  ○ Histopathology analysis
  ○ single-cell RNA sequencing (scRNA-seq) analysis
  ○ Quantitative real-Time (qRT-PCR) validation
  ○ Amino acid uptake assay
  ○ Amino acid uptake in T cells under small-molecule treatments
  ○ Amino acid uptake in Th17-polarized T cells under small-molecule treatments
  ○ Statistical Analysis and reproducibility
- ADDITIONAL RESOURCES

## STAR ★ METHODS

KEY RESOURCES TABLE.

2 KEY RESOURCES TABLE
| REAGENT OR RESOURCE | SOURCE | IDENTIFIER |
| --- | --- | --- |
| **Antibodies** |  |  |
| *Brucella* polyclonal antibody | Bioss Antibodies Inc. | bs-2229R |
| Goat Anti-Rabbit CF 560 Antibody | Biotium | Cat# 20300-1 |
| anti- Ly6g antibody | Biolegend | Cat# 127605 |
| anti-CD45-violetFluor™ | Cytek Biosciences | Cat# 85-0454 |
| anti-CD3-APC/Cy7 | Biolegend | 17A2 |
| anti-CD4-Alexa Fluor 700 | Biolegend | RM4-4 |
| anti-CD8a-BV570™ | Biolegend | 53-6.7 |
| anti-CD19-PE/Dazzle™ 594 | Biolegend | 6D5 |
| anti-CD11c-APC | Biolegend | N418 |
| anti-CD11b- BV750™ | Biolegend | M1/70 |
| anti-CD163-APC/ Fire™ 810 | Biolegend | S15409I |
| anti-NK1.1-BV785™ | Biolegend | PK136 |
| anti-FoxP3-Pacific Blue™ | Biolegend | MF-14 |
| anti-CD25- BV605™ | Biolegend | PC61 |
| anti-CD69-PerCP/Cy5.5 | Biolegend | H1.2F3 |
| anti-Ly6C-PE/Cy7 | Biolegend | HK1.4 |
| anti-Ly6G- BV650™ | Biolegend | 1A8 |
| anti-PD-1- BV421™ | Biolegend | RMP1-30 |
| anti-MHC-II- cFluor® BYG750 | Cytek Biosciences | M5/114.15.2 |
| anti-F4/80- Alexa Fluor 647 | Biolegend | QA17A29 |
| anti-Zap70- PE | Biolegend | 1E7.2 |
| anti-Pax5-Alex Fluor 488 | Biolegend | 1H9 |
| anti-FcgRIV- BV711™ | Biolegend | 9.00E+09 |
| Mouse anti-PD-1 | Bio X cell | RMP1-14 |
| anti-TNF-α | Biolegend | MP6-XT22 |
| anti-perforin | Biolegend | Cat# 154404 |
| anti-granzyme B (Gr-B) | Biolegend | GB11 |
| Purified anti-mouse CD3 antibody (17A2) | Biolegend | Cat# 100202 |
| Purified anti-mouse CD28 antibody (37.51) | Biolegend | Cat# 102102 |
| anti Ki-67 |  |  |
| Anti-CD90.2 (THY1) microbeads | Miltenyi Biotec | Cat# 130-121-278 |
| **Biological samples** |  |  |
| Bone marrow-derived macrophages (BMDMs) | This study |  |
| B16-Ova tumor samples | This study |  |
| **Chemicals, peptides, and recombinant proteins** |  |  |
| Trizol | Invitrogen | 15596026 |
| Superscript™ III First-Strand Synthesis | Invitrogen | 18091050 |
| SYBR Green | Invitrogen | 11761100 |
| Tryptone Soy Broth (SOB) | Millipore Sigma, USA |  |
| DMEM medium | Invitrogen, CA, USA |  |
| RPMI 1640 medium | Invitrogen, CA, USA |  |
| Penicillin-streptomycin solution, 100× | Millipore Sigma, USA |  |
| Tryptic Soy Agar (TSA) | Millipore Sigma, USA |  |
| Chloramphenicol | Research product international, IL, USA |  |
| D-Luciferin | Promega |  |
| Sodium borate buffer |  |  |
| ACK lysis buffer | Gibco, USA | Cat # A1049201 |
| Mouse GM-CSF | PeproTech | Cat # 315-03-5UG |
| 6-aminoquinolyl-N-hydroxysuccinimidyl carbamate | Cayman Chemical, MI, USA |  |
| Advanced RPMI 1640 Medium | Gibco | Cat #12633012 |
| FBS | Gibco | Cat # 10-082-147 |
| IL-2 | Stemcell Tech. | Cat # 78081 |
| CD3/CD28 dynabeads | Thermo | Cat # 11456D |
| Indole | Sigma | Cat# I3408 |
| 5-hydroxyindole | Sigma | Cat# H31859 |
| BAY | MedChemExpress | Cat# HY-135829 |
| FICZ | Sigma | Cat# SML1489 |
| DMSO (control solvent) | Sigma |  |
| Recombinant IL-6 | STEMCELL Technologies | Cat# 78052.1 |
| TGF-β | BioLegend | Cat# 763102 |
| **Critical commercial assays** |  |  |
| CD8a+ T Cell Isolation Kit, mouse | Biolegend | Cat# 480008 |
| Zombie Aqua™ Fixable Viability Kit | Biolegend | Cat# 423101 |
| MojoSort™ Mouse CD4 T Cell Isolation Kt | Biolegend | Cat# 480006 |
| MACS Tumor Dissociation Kit, mouse | Miltenyi Biotek | Cat# 130-096-730 |
| Aqua Zombie Fixable Dead/Live Staining Kit | BioLegend | Cat# 423101 |
| CFSE staining kits |  |  |
| **Deposited data** |  |  |
| scRNA-seq data | This paper | GEO: XXXXX |
| Metabolomics data | This paper |  |
| **Experimental models** |  |  |
| **Bacteria strains** |  |  |
| SPIKE chassis | This paper |  |
| SPIKE 1.0 | This paper |  |
| **Cell lines** |  |  |
| H1975 | ATCC | ATCC CRL 5908, RRID: CVCL_1511 |
| Panc02 | ATCC | ATCC CRL 2553, RRID: CVCL_D627 |
| AsPC-1 | ATCC | ATCC CRL 1682, RRID: CVCL_0152 |
| B16-Ova |  |  |
| LLC1-OVA | Vitro Biotech | VOA1B004 |
| KPC-luciferase-expressing cells | This paper | Gift from Dr Craig Logsdon,<br>MD Anderson Cancer Center |
| **Mouse lines** |  |  |
| C57BL/6J WT | The Jackson Lab | RRID: IMSR_JAX: 000664 |
| OT-1 | The Jackson Lab | RRID: IMSR_JAX: 003831 |
| NSG | The Jackson Lab | RRID: IMSR_JAX: 005557 |
| CD34<sup>+</sup> humanized mouse | The Jackson Lab |  |
| CD1d KO mice | The Jackson Lab | RRID: IMSR_JAX:008881 |
| CD4 KO mice | The Jackson Lab | RRID: IMSR_JAX:002663 |
| CD8 KO mice | The Jackson Lab | RRID: IMSR_JAX: 002665 |
| Batf3 KO mice | The Jackson Lab | RRID: IMSR_JAX: 013755 |
| AhR-KO macrophages |  |  |
| **Ruminant Animals** |  |  |
| Goats |  | *Capra hircus* outbred,<br>pregnant |
| **Oligonucleotides** |  |  |
| See Table S5 for qPCR primers |  |  |
| **Software and algorithms** |  |  |
| FlowJo v10.9.0 | FlowJo LLC | https://www.flowjo.com/ |
| FlowJo version 10.10 | Tree Star Inc. OR, USA |  |
| GraphPad-Prism v10.2.2 | Graphpad Software LLC | https://www.graphpad.com |
| Living Image v4.5.5 | PerkinElmer | https://www.perkinelmer.com/ |
| Quantum GX2 software with 3D viewer | PerkinElmer | https://www.perkinelmer.com/ |
| Leica Application Suite X | Leica | https://www.leica-microsystems.com |
| Aperio ImageScope v12.3.3 | Leica | https://www.leica-microsystems.com |
| ImageJ | National Institute of Health | https://imagej.net/ij/ |
| Cell Ranger 7.1.0 pipeline |  |  |
| Living Image 4.8.0 | Revvity Health Sciences Inc., USA | https://www.revvity.com/software-downloads/in-vivo-imaging |
| QuPath software | Bankhead et al. 62 | https://qupath.github.io/ |
| MetaboAnalyst |  | https://metaboanalyst.ca/ |
| Seurat V4 | Hao et al. 63 | https://satijalab.org/seurat/articles/get_started.html |
| PanglaoDB | Franzen et al. 64 | https://panglaodb.se/ |
| AUCell | Aibar et al. 65 | https://www.bioconductor.org/packages/release/bioc/html/AUCell.html |
| **Other** |  |  |
| None |  |  |

## METHOD DETAILS

### Bacterial culture and metabolic engineering

Bacterial strains SC and S1.0 (Key Resources Table) was cultured as previously described^66–68^. For SC or S1.0 administration, tryptone Soy Broth (SOB) (Millipore Sigma, USA) was used to culture the bacterial strains. The bacteria were collected by centrifugation followed by washing and resuspending in 1 × PBS. For co-culture bacteria with cells, bacteria were added to each well of a 24-well plate at an MOI (Multiplicity of infection) of 20 of macrophage monolayer in DMEM medium (Invitrogen, CA, USA).

### Generation of HI-producing S1.0

To generate a SPIKE strain that produces HIs, we cloned *tna* and *tmo* encoding tryptophanase (Tna) and toluene-4 monooxygenase (Tmo), respectively, into a broad range bacteria expression plasmid pBBR1MCS6Y^69^ and transferred the plasmid to BmΔ*vjbR*^6,14^. Briefly, *Brucella* codon-optimized *tna* (for Tna) from *Escherichia* coli K-12^14^ and *tmo* (for Tmo A-F) from *Pseudomonas mendocina*^70^ were cloned into the *Brucella*-preferred, broad-host-range plasmid pBBR6Y, a pBBR1MCS derivative, under the control of J23119 and pAphA promoters. The resulting plasmid was then introduced into SC to generate strain S1.0.

### Cell culture

All mammalian cells used in this work are listed in Key Resources Table. H1975 human non-small cell lung cancer cells, Panc02 mouse pancreatic ductal adenocarcinoma cells, and AsPC-1 (human pancreatic cancer cells derived from nude mouse xenografts) cells were cultured in 10% FBS, 1% penicillin-streptomycin supplemented RPMI 1640 medium (Invitrogen, CA, USA). B16-Ova melanoma cells were grown in DMEM medium supplemented with 10% FBS and 1% penicillin-streptomycin antibiotics.

### Human CD8^+^ T cell isolation and characterization

Human CD8⁺ T cells were isolated from de-identified blood bags obtained from healthy donors (n = 15 independent blood bags). Purified CD8⁺ T cells were cultured in complete RPMI 1640 medium and activated using anti-CD3 and anti-CD28 antibodies. Briefly, plates were coated with anti-CD3 antibody (e.g., 1–5 µg/mL), and soluble anti-CD28 antibody (e.g., 1–2 µg/mL) was added at the time of stimulation. Cells were cultured under standard conditions (37 °C, 5% CO₂) and treated with commercial 5-hydroxyindole (5-HI) as the indicated concentration (50 μM) determined by the dose-response assay (Figure S1A). Following stimulation, cells were harvested and analyzed by flow cytometry for activation and functional markers.

### Flow cytometric analysis of cytokine production in CD3/CD28-stimulated CD8⁺ and CD4^+^ T cells following 5-HI treatment

Purified murine CD8⁺ T cells and/or CD4^+^ T cells were activated using plate-bound anti-CD3 antibody (2 µg/mL) in combination with soluble anti-CD28 antibody (1 µg/mL), as above-described. Cells were cultured under non-polarizing conditions, and no additional cytokines were added for directed CD4⁺ T helper differentiation. Following stimulation, cells were harvested and analyzed by flow cytometry for activation markers. Human CD8^+^ T cells were isolated from freshly derived PBMCs from blood and activated using human anti CD3/CD28 antibodies. All the T cell isolation were performed by negative selection and magnetic separation using the MojoSort Mouse or Human CD8 T Cell Isolation Kit (BioLegend, USA). At 3 days post-activation (dpa), the cells were treated with 50 µM of commercial 5-HI or indole (the relevant metabolic precursor with immunomodulatory and anti-inflammatory effects^14^) or vehicle control. The levels of TNF-*a*, Perforin, and GrB (for murine CD8⁺, CD4^+^ T cells) and TNF-*a* and IL-2 – were tested at 48 hours post treatment by flowcytometric analysis as previously described^6^. The same protocol was followed for assessing CD4^+^ T cells differentiation in Th17 cells.

### Isolation and characterization of bone marrow derived macrophages (BMDMs)

Murine BMDMs were generated as previously described with minor modifications^71,72^. Briefly, bone marrow cells were harvested from the femurs and tibias of adult mice under sterile conditions, and red blood cells were lysed using ACK lysis buffer (Gibco USA). Cells were cultured in complete RPMI 1640 medium supplemented with 10% fetal bovine serum, 1% penicillin– streptomycin, 2 mM L-glutamine, and recombinant murine granulocyte–macrophage colony-stimulating factor (GM-CSF; 20 ng/mL) for 7 days to allow macrophage differentiation. Fresh medium containing GM-CSF was replenished on days 3 and 5. Differentiated macrophages were characterized by flow cytometry, and M1 polarization was confirmed based on CD38 expression.

### *In-vitro* cytotoxic assessment of CD8^+^ T cells activity against various cancer cells

BMDMs were isolated from C57BL/6 mouse (n=15) and expanded with GM-CSF and were used as antigen-presenting cells to prime antigen-specific CD8⁺ T cells. Subsequently, these BMDMs were co-cultured with S1.0, SC at an MOI of 20 or remained untreated in the presence of antigen-specific CD8^+^ T cells as previously reported^6^. The CD8^+^ T cells were then retrieved from the culture and added to the target B16-Ova or H1975 cancer cells at various target to effector ratios of 1:1, 1:2 and 1:5. Dead/live staining was performed using the Aqua Zombie Fixable Dead/Live Staining Kit (BioLegend, Cat# 423101) on CFSE-pre-stained CD8^+^ T cells to distinguish them from cancer cells and assess cancer cell death.

### Identification of HIs from S1.0 bacterial culture filtrate

Preparation of S1.0 bacterial culture filtrate performed as previously described^73^. Briefly, S.10 bacteria were cultured in SOB for 48 hours, culture media were collected and centrifuge at ∼20,160 ×g for 15 min to remove cell debris, media were then separated by methanol-chloroform-mQ water extraction, the upper (aqueous) phase was transfer into an Eppendorf tube and dried using a SpeedVac vacuum concentrator (Thermo Fisher Scientific Inc., USA). Detection and identification of HIs were performed using Liquid Chromatography-Mass Spectrometry (LC-MS) under established conditions. These included defined chromatographic separation (mobile phase composition and gradient), flow rate, and mass spectrometry acquisition parameters optimized for HI detection, as described previously^73^. Based on mass spectrometry identification of multiple HIs, 5-HI was selected as a representative metabolite for all *ex vivo* stimulation assays in this work after dose-response evaluation of its bioactivity, as it was sufficient to reproduce the inflammatory effects.

### Mouse lines

All the mouse lines used in this study and their origins are listed in Key Resources Table. The wild-type C57BL/6 (B6) Thy 1.1 mice were purchased from the Jackson Laboratories. Non-obese diabetic-Severe combined immunocompromised γ (NOD.Cg-Prkdcˢᶜⁱᵈ Il2rgᵗᵐ¹ᵂʲˡ/SzJ; NSG) mouse model mice are severely immunocompromised mice as they lack the IL-2 receptor γ chain. The mice were procured from Jackson laboratories and maintained in 12 h light and dark cycles with their littermates. The humanized mice (huCD34 mice) are created by transferring CD34^+^ hematopoietic stem cells into NSG mice to mimic a humanized immune system. The humanized mice were procured from Charles River laboratories and maintained under similar conditions. The OT-1 transgenic mice produce a transgenic T-cell receptor on CD8^+^ T cells recognizing ovalbumin peptide residues 257-264 (OVA_257-264_). OT-1 T cell receptor transgenic mice were procured from Jackson Laboratories. Six to eight weeks old mice, unless otherwise indicated, were used in each specific mouse tumor model described details as the below.

### S1.0 safety assays

#### Mouse complete blood count (CBC) assays

C57BL/6 mouse organ weight analysis was conducted on mice intravenously injected with varying doses of S1.0 (n=5 per group), receiving either 100 µl of the S1.0 bacterial dose or 100 µl of 1× phosphate-buffered saline (PBS). Organ (spleen, liver, lung, kidney, brain) weights were collected on 14 days post-injection (DPI). Additionally, a CBC analysis, including red cell distribution width, neutrophil count, reticulocyte absolute count (Retic Count), lymphocyte count, hematocrit, hemoglobin, red blood cell count, white blood cell count, mean corpuscular volume, mean corpuscular hemoglobin mean corpuscular hemoglobin concentration (MCHC), and plasma protein, was performed on mice intravenously administered with different S1.0 bacterial doses or 1× PBS. Results were analyzed on 14 DPI.

#### S1.0 biodistribution in tumor-bearing mice

Male C57BL/6 wild type mice were subcutaneously injected with 1×10^6^ LLC1-Ova cells (Vitro Biotech, CA, USA) (Key Resources Table) in the right lateral flank on day 0. The mice were divided into three groups (n=15 per group). On day 8, once the tumor volume reached approximately 150 mm^3^, mice were intravenously injected with 5.0 × 10^7^ S1.0 (group 1), 1.0 × 10^8^ S1.0 (group 2), 1.0 × 10^9^ S1.0 (group 3), and 5.0 × 10^7^ S1.0 + CAR-CD8 T cells (group 4), respectively. Intravenous delivery of S1.0 enables systemic distribution and access to tumor sites. For the group 4, tumor-bearing mice were injected intravenously with 2 × 10^6^ murine CAR-CD8 T cells in combination with 5.0 × 10^7^ S1.0 bacteria. The colony forming unit (CFU) assay was performed to determined S1.0 bacterial distribution in tumor and other organs of mice pre-implanted with or without tumor cells and injected with S1.0 as a tumor therapy treatment as described previously^14^. In brief, mice were humanely sacrificed using CO₂ inhalation followed by cervical dislocation at 4-, 8-, 12-, 16-, and 20-DPI, and tumor, spleen, lungs, kidney, liver, brain, bone marrow, fecal and urine samples were collected. The harvested organs were homogenized and plated on Tryptic Soy Agar (TSA) (Millipore Sigma, USA) plates supplemented with chloramphenicol (30 μg/mL) (Research product international, IL, USA). The CFU was recorded after 3-4 days of post-cultivation of the bacteria. The CFU was calculated using the following formula: CFU/g = (Number of colonies*dilution factor) / weight of tissue sample in gm.

#### Goat S1.0 safety assessment

Two groups of pregnant goats (n=7/group) at approximately 74 days of gestation were inoculated bilaterally in the conjunctival sac with a total dosage of 1×10^8^ and 1×10^9^ S1.0 bacteria, respectively. Clinical evaluations were conducted daily, and blood samples were collected bi-weekly for card test and ELISA analysis through 84 days post S1.0 inoculation and at euthanasia. The dams had normal parturitions and healthy kids. At parturition, colostrum, vaginal swab, placenta, umbilical cord, and urine were collected, and following humane euthanasia at approximately 10 days post-parturition, multiple tissues were collected from the dam and kids for histology, bacteriology, and quantitative PCR to detect DNA from S1.0 using a S1.0-specific probes (IS711-FAM, and tmo-Cy5) (Table S5).

### CAR-T and TCR-T cell preparation

The MSGV1 γ retroviral vector backbone was modified to express mesothelin antigen specific scFv as described in our previous studies^75,76^. CAR-T cells were prepared by transduction of viral supernatants containing the respective antigen with 5 µg/mL polybrene. The lentivirus to modify NY-ESO-TCR specific CD8^+^ T cells derived from human blood peripheral blood mononuclear cell sample – MSGV1 γ retroviral vector backbone was kindly provided by Dr. Wendell Yang (Imgen Lifesciences Inc.).

### Murine tumor models

Tumor cell inoculation and cancer treatments using divergent murine tumor models are illustrated in Figure 2A-B, Figure 3A, and Figure 5A. Six to eight weeks old mice (Key Resources Table) were used in each specific mouse tumor model described details as the below.

#### Lung cancer model in NSG mice and hCD34^+^ humanized mice

Wild-type NSG mice or hCD34^+^ NSG humanized mice were subcutaneously injected with 1×10^6^ H1975 human lung cancer cells in the right lateral flank on day 0. The mice were then divided into five different groups (n = 5 mice/ per group). The first 3 groups of mice were injected intravenously with either 1× PBS control (untreated control), live attenuated 1 × 10^7^ SC bacteria or engineered live attenuated S1.0 bacteria on day 7 post tumor-cell inoculation (PTCI). On Day 9, two more groups of mice were injected intravenously with 2 × 10^6^ NY-ESO-TCR modified CD8^+^ T cells or NY-ESO-TCR modified CD8^+^ T cells in combination with 1 × 10^7^ S1.0 bacteria. Mice were housed in Texas A&M University, Laboratory Animal Resources and Research Facility, and health status checked daily. The survival of tumor bearing mice was monitored over 60 days. Tumor growth was monitored every other day and tumor volumes were calculated using the formula: Tumor Volume (mm^3^) =0.5 × length × width^2^^75,77^. Mice were humanely euthanized if tumor size reached above 4000 mm^3^.

#### MC32-CEA Colon cancer, Panc02 pancreatic cancer and B16 melanoma models in C57BL/6 mice

Wild-type C57BL/6 (B6) Thy 1.1 mice (Jackson Laboratories) were subcutaneously injected with 1×10^6^ MC32-CEA colon cancer cells, Panc02 mouse pancreatic cancer cells, or B16-ovalbumin melanoma cancer cells in the right lateral flank on day 0. For each mouse tumor model, the mice were then divided into five different groups (n = 5 mice per group). The first 3 groups of mice were injected intravenously with either 1× PBS control, 1× 10^7^ SC or S1.0 bacteria on day 7 PTCI. For MC32-CEA colon cancer model, on Day 9, two more groups of mice were injected intravenously with 2 × 10^6^ CAR-CEA modified CD8^+^ T cells only or CAR-CEA modified CD8^+^ T cells in combination with S1.0 bacteria. For Panc02 pancreatic cancer model, on Day 9, two more groups of mice were intravenously injected with 2 × 10^6^ mesothelin-modified CAR-CD8 T cells only or mesothelin-modified CAR-CEA modified CD8^+^ T cells in combination with 1 × 10^7^ S1.0 bacteria. For B16-Ova melanoma cancer model, on Day 9, two more groups of mice were intravenously injected with 2 × 10^6^ OT-1 CD8 T cells only or OT-1 CD8 T cells in combination with 1 × 10^7^ S1.0 bacteria. Housing and checking health status of the treated mice, monitoring tumor growth and survival of tumor-bearing mice were performed as the methods described above.

#### Pancreatic cancer model in NSG mice

NSG mice were subcutaneously injected with 1×10^6^ AsPC-1 human pancreatic cancer cells in the right lateral flank on day 0. The mice were then divided into five different groups (n=5 mice per group). The first 3 groups of mice were intravenously injected with either 1× PBS control, 1 × 10^7^ SC or S1.0 bacteria on day 7 PTCI. On Day 9, two more groups of mice were injected intravenously with 2 × 10^6^ mesothelin-modified CAR-CD8 T cells in combination with 1 × 10^7^ S1.0 bacteria. Housing and checking health status of the treated mice, monitoring tumor growth and survival of tumor-bearing mice were performed as the methods described above.

#### KPC-Luc PDAC and B16F10-Luc melanoma tumor in gene knockout (KO) mice

CD1, CD4, CD8, or Batf3 KO and control mice were intraperitoneally injected with 1×10^6^ KPC-Luc or B16F10-Luc melanoma tumor cells on day 0. For each tumor gene KO or control mouse model, the mice were divided into four two groups (n=10 mice per group). On day 14 PTCI, the two groups of mice were intraperitoneally injected with 100 μl 1× PBS or 1 × 10^7^ S1.0 bacteria in PBS. Mice were housed in the University of Missouri, Laboratory Animal Resources and Research Facility. Checking health status of the treated mice, monitoring tumor growth and survival of tumor-bearing mice were performed as the methods described above.

#### Orthotopic tumor model and treatment with S1.0 and antibodies to co-inhibitory molecules

Six-week-old female albino C57BL/6 were obtained from The Jackson Laboratory (Bar Harbor, ME). Orthotopic pancreatic ductal adenocarcinoma was established by surgically implanting Kras^G12D^; P53^flox/flox^; PDX-1-Cre; Luciferase [KPC-Luc] cells into the pancreas of 7-8 week old mice (n=10 mice/group) as described previously^78^. Establishment of tumors was confirmed 7 days post-surgical implantation using IVIS (in vivo imaging system) bioluminescence imaging (IVIS Spectrum, Revvity, Waltham, MA) and mice were randomized for different treatment groups. Tumor bearing mice were treated with either S1.0 (5.0 × 10^7^ CFU/mouse) administered intravenously or 200 µg/mouse of anti-PD1 (αPD-1, clone: RMP1-14, BioXCell, Lebanon, NH) intraperitoneally, or a combination of αPD-1 and S1.0. Control mice received manufacture recommended rat Ig2a isotype control antibody (BioXCell, Lebanon, NH). Mice were treated with αPD-1 or isotype antibody, intraperitoneally, on days 8, 10, 13, 19, 22, and 25 (Figure 3A). IVIS bioluminescence imaging with 150 mg/kg body weight (BW) of D-Luciferin (Promega) administered intraperitoneally was used to monitor tumor growth two times a week for survival studies. To quantify tumor growth, regions of interest were manually drawn around tumor borders using Living Image software (Living Image 4.8.0, Revvity Health Sciences Inc., USA) and average tumor radiance (p/s/cm^2^/sr) was determined. For determining the infiltration of immune cells in the tumor with different treatments, mice were humanely sacrificed on days 15 PTCI (Figure 3A), tumor tissues and spleen were collected, and single-cell suspensions were prepared for flow cytometry analysis^81^. All animal procedures were reviewed and approved by the MD Anderson Cancer Center Animal Care and Use Committee.

### Fluorescent cytometric analysis of biomarker expression in PDAC tumor-bearing mice

Immune cells isolated from tumor and spleen of PDAC tumor-bearing mice were stained with a panel of anti-mouse antibodies (Key Resources Table) that included anti-CD45-violetFluor^TM^ (104, Cytek Biosciences, CA, US), anti-CD3-APC/Cy7 (17A2), anti-CD4-Alexa Fluor 700 (RM4-4), anti-CD8a-BV570^TM^ (53-6.7), anti-CD19-PE/Dazzle^TM^ 594 (6D5), anti-CD11c-APC (N418), anti-CD11b-BV750^TM^ (M1/70), anti-CD163-APC/ Fire^TM^ 810 (S15409I), anti-NK1.1-BV785^TM^ (PK136), anti-FoxP3-Pacific Blue^TM^ (MF-14) anti-CD25-BV605^TM^ (PC61), anti-CD69-PerCP/Cy5.5 (H1.2F3), anti-Ly6C-PE/Cy7 (HK1.4), anti-Ly6G-BV650^TM^ (1A8), anti-PD-1-BV421^TM^ (RMP1-30), anti-MHC-II-cFluor® BYG750 (M5/114.15.2, Cytek Biosciences, CA, US), anti-F4/80-Alexa Fluor 647 (QA17A29), anti-Zap70-PE (1E7.2) anti-Pax5-Alex Fluor 488 (1H9), and anti-FcgRIV-BV711^TM^ (9E9), which were purchased from Biolegend (San Diego, CA, USA) unless otherwise indicated. One to two million cells were stained in a volume of 100 μl and the concentration of each antibody in the cocktail was 0.05 μg/sample. The data were acquired on Cytek Northern Lights spectral flow cytometer (Cytek Biosciences, CA, US) and analyzed using FlowJo version 10.10 (Tree Star Inc. OR, USA).

### Spatial multiplex immunohistochemical imaging analysis

Single cell recognition and quantitative biomarker analysis were performed on subcutaneous Panc02 mouse pancreatic cancer tissues, dissected from tumor-bearing C57BL/6 mice (n=3) by using Canopy Biosciences’ ChipCytometry™ platform with the CellScape™ instrument (Canopy Biosciences; a Brucker Company). A 14-plex assay panel was performed on four explanted tumor samples from the groups of 1) control (untreated sample), 2) CAR-T cell only, 3) S1.0, and 4) S1.0 combination with CAR-T cells. Gating and quantification of target cell population were performed, and the percentages and cell counts were determined using the CellScape™ system. Single cell segmentation was performed on high-resolution images allowing for the phenotyping of each individual cell by cell classification/thresholding. For further spatial analysis, QuPath software^62^ was utilized to segment the images into equal-sized regions to calculate cell densities for CD11b+ monocytes, CD8+ T cells, CD4+ T cells, and their co-expression with PD-1 and Ki-67.

### Multiphoton image analysis

Multiphoton microscopy was conducted from the Day 21 explanted B16-Ova melanoma tumors by using an anti-*Brucella* rabbit polyclonal antibody (bs-2229R; Bioss Antibodies Inc.) and secondary anti-rabbit CF 560 Antibody (Cat# 20300-1; Biotium). MDSCs staining was performed by using the anti-Ly-6g antibody (Biolegend). Antibodies used in this work are listed in (Key Resources Table).

### Metabolomics analysis

Metabolites were isolated from explanted tumor tissues of untreated control, SC and S1.0 treated B16-Ova melanoma tumor-bearing wildtype C57BL/6 mice (n=5 per treatment group) at day 21 PTCI. The polar solvents were then dried and analysed by mass spectrometry analysis. For each metabolite, the average expression values across three replicates of each treatment condition were calculated. PLS-DA was performed by using the metabolomics analysis software MetaboAnalyst (https://metaboanalyst.ca/) to perform peak alignment and identification. Then, hierarchical clustering analysis was done on the z-normalized average expression values of metabolites to group the data based on their expression patterns across tumor conditions. The heatmap of metabolites group with higher expression values in the S1.0 treated tumours compared to the controls is plotted.

### Histopathology analysis

After euthanasia, representative samples of the tumor mass from each mouse were collected and fixed by immersion in 10% neutral buffered formalin at room temperature for 24 hours and then stored in 70% ethanol before embedding in paraffin, sectioning at 5 μm and staining with hematoxylin and eosin (H&E). Bright field whole slide images of H&E-stained tissue sections were captured as digital files by scanning at 20 × using a 3DHistech Panoramic SCAN II FL scanner (Budapest, Hungary). The scanned images were evaluated by a board-certified veterinary pathologist in a blinded manner and scored on a scale from 0 = normal to 4 = extensive for local tissue invasion and abundance of lymphocytes, neutrophils, macrophages and necrosis^82,83^. Statistical analysis of the histopathology ordinal data was performed by using the GraphPad Prism software (Dotmatics, https://www.graphpad.com)

### single-cell RNA sequencing (scRNA-seq) analysis

scRNA-seq analysis was performed on tumor tissues derived from B16-Ova melanoma tumor-bearing wildtype C57BL/6 mice (n=5 per treatment group). The tumor was dissociated by using MACS tumor dissociation kit (Miltenyi biotec) and viable single cell suspension were analysed and run on 10× Genomics platform. Raw sequencing reads from single-cell RNA sequencing (scRNA-seq) data were processed using the Cell Ranger 7.1.0 pipeline (10× Genomics). Reads were aligned to the mm10 reference genome, and gene expression matrices were generated. All groups were pooled together and then QC filtered and preprocessed together to control bias. scRNA-seq data analysis was performed on R using the Seurat v4^63^. Cells with < 500 reads and expressing > 15 percent mitochondrial genes expression were filtered out^84^. Genes expressing in < 3 cells were filtered out. The filtered data were then library size normalized and scaled to 10000 total, followed by log(X+1) transformation using the ‘NormalizeData’ function. The top 2000 highly variable genes (HVG) were selected using ‘FindVariableFeatures’ function for principal component analysis (PCA). Uniform Manifold Approximation and Projection (UMAP) was performed using the top 15 principal components to project the data onto a two-dimensional space for visualization. Cells were then clustered using the Louvain algorithm with a resolution of 0.8 and 22 cell clusters were identified.

#### Annotation

Significant marker genes for each cluster (Table S1) were identified by performing differential analysis using the Wilcoxon rank sum test using the ‘FindAllMarkers’ function. Each individual cluster was annotated using the marker genes and canonical cell type markers from PanglaoDB^64^.

#### Differential expression, enrichment analysis and cell scoring

Differentially expressed gene (DEG) analysis was performed using the Wilcoxon rank-sum test for each cell type and across all four treatment conditions. Significant DEGs were selected by an adjusted P value cut-off of 0.05. Top 100 DEGs for each cell type were used for gene list enrichment analysis with the KEGG_2019_Mouse database using Enrichr^85^. Cell scores for gene sets were calculated using the AUCell^65^ for gene sets. Gene sets obtained from the KEGG Mouse Pathway Database and T-cell exhaustion markers were retrieved as previously described^86^. Gene expression and cell proportion were visualized in R using Seuratv4’s inbuilt plotting functions, ggplot2 and dittoSeq.

### Quantitative real-Time (qRT-PCR) validation

Total RNA was isolated from fresh control and S1.0-treated and B16-Ova melanoma tumor tissue homogenate using Trizol (Invitrogen cat no 15596026) and reverse transcribed into cDNA using the Superscript™ III First-Strand Synthesis system (Invitrogen Cat# 18091050) as reported previously^87^. Gene expression was determined by qRT-PCR using the CFX duet Real-Time PCR Detection System (Bio-Rad, USA) and SYBR Green (Invitrogen Cat# 11761100). The RT-PCR was performed to determine the listed genes using the mouse-specific primers (Table S5). Melt curves were generated, and fold expression was calculated by using the 2^−ΔΔCt^ method, which was normalized using β-actin as the endogenous control as described previously^87^.

### Amino acid uptake assay

The tumors from tumor-bearing mice of various experimental groups were weighed, chopped, and placed into gentleMACS M Tubes (Miltenyi Biotec, Cat# 130-093-236). A mixture of methanol, acetonitrile, and water (5:3:2, v/v/v) was added at a volume equivalent to five times the tumor weight, and the samples were homogenized using the gentleMACS^TM^ Dissociator (Miltenyi Biotec). The homogenate was then transferred to 1.5 ml Eppendorf tubes and centrifuged at 13,000 × g for 15 minutes. The resulting supernatant was collected in glass vials and dried under a gentle stream of nitrogen gas. The dried extracts were reconstituted in 40 μl of water/acetonitrile (8/2, v/v), and 10 μl of the solution was transferred to a glass autosampler vial, followed by the addition of 35 μl sodium borate buffer (100 mM, pH 9.0). Subsequently, 10 μl of 6-aminoquinolyl-N-hydroxysuccinimidyl carbamate (AQC, 10 mM in acetonitrile) derivatizing reagent (Cayman Chemical, MI, USA) was added. The vials were tightly capped, vortexed, and incubated at 55 °C for 15 minutes. After cooling to room temperature, 1 μL of the derivatized sample was analyzed by liquid chromatography-tandem mass spectrometry (LC-MS/MS). LC separations were performed using an ACQUITY Premier UPLC System (Waters Corporation, MA USA) equipped with a C_18_ ACQUITY UPLC BEH Column (2.1 mm × 150 mm, 1.7 μm, 130 Å, part no: 186002353) operated at 50 °C. The mobile phase consisted of water with 0.1% formic acid (mobile phase A) and acetonitrile (mobile phase B), with a non-linear gradient program as follows: 0−2 min, 3% B; 2−8 min, 3–40% B; 8−8.1 min, 96% B; 8.1−12 min, 96% B; 12−12.5 min, 3% B; 12.5−14 min, 3% B. The flow rate was 500 μL/min, and the injection volume was 1 μL. Detection was performed using a Xevo-XS Triple Quadrupole Mass Spectrometer (TQ-XS) (Waters Corporation, MA USA) operated in positive ion mode with multiple reaction monitoring (MRM) mass spectrometry^88^.

### Amino acid uptake in T cells under small-molecule treatments

Single-cell spleen suspensions were prepared from C57BL/6 mice, and total T cells were isolated using anti-CD90.2 microbeads (Miltenyi Biotec. Cat# 130-121-278) following the manufacturer’s protocol. Purified T cells were counted, cultured in Advanced RPMI 1640 Medium (Gibco, Cat# 12633012) in presence of 10% FBS (Gibco, Cat# 10-082-147) and IL-2 (Stemcell Tech. Cat# 78081) and activated using CD3/CD28 dynabeads (Thermo, Cat# 11456D) for 3 days according to manufacturer’s protocol. After polarization, cells were split into treatment groups and exposed to the indicated small molecules for 24h: Indole (50 µM, Sigma, Cat# I3408), 5-hydroxyindole (5-HI; 50 µM, Sigma, Cat# H31859), BAY (1 µM, MedChemExpress, Cat# HY-135829)^44^, or FICZ (100 nM, Sigma, Cat # SML1489)^89^; a control (matching solvent) was included in each experiment. At 24 h after initiating treatment, cells were washed and subjected to the amino-acid uptake assay by spiking cultures with the appropriate isotope-labeled amino acid for 30 min. Uptake was quantified as previously described^34^ and normalized to viable cell counts measured immediately prior to the labeling pulse exactly as described in “Amino acid uptake assay”.

### Amino acid uptake in Th17-polarized T cells under small-molecule treatments

Single-cell spleen suspensions were prepared from C57BL/6 mice, and total T cells were isolated and purified as the above-mentioned methods but under Th17-polarizing conditions for 3 days with recombinant IL-6 (20 ng/mL; Stemcell Tech. Cat# 78052.1) and TGF-β (1 ng/mL; BioLegend, Cat# 763102). After polarization, cells were split into treatment groups and exposed to the indicated small molecules for 24h as described above. At 24 h after initiating treatment, cells were washed and subjected to AA uptake assay as describe above.

### Statistical Analysis and reproducibility

The statistical differences between two groups and among groups were determined using Student’s t-test and one-way ANOVA, respectively. Tumor growth curves were compared using a two-way ANOVA. Tukey correction was used for multiple comparisons test. Statistical significance for survival analysis was analyzed using the Kaplan–Meier method and compared among groups using the Wilcoxon and log-rank test when indicated. All tests were two-tailed with a considered significance level of *p* < 0.05. All the statistical analysis were calculated using Graphpad Prism version11 (GraphPad Software). Graphs show the mean ± standard error of the means (SEM) unless otherwise indicated. All *in vitro* experiments were independently repeated at least three times. All *in vivo* mouse experiments were conducted with 5 mice per group (n=5/group) unless otherwise indicated and were repeated at least twice. The *in vivo* goat safety experiment was performed with groups of pregnant does (n=7/group) for each of the two doses. All the *in vitro* and *in vivo* independently repeated experiments produced consistent or similar results.

### Extended Data Figures and Figure legends

**Figure S1.** Hydroxyindole (HI) increases cytokine production and cytotoxicity of CD8^+^ T cells.

**Figure S2.** Safety assessment of SPIKE 1.0 (S1.0) in mice and pregnant goat models.

**Figure S3.** Chip-cytometry multiplex imaging analysis of biomarker expression in the TME of B16-Ova melanoma tumor-bearing mice at day 28 PTCI.

**Figure S4.** Identification of different cell types by scRNA-seq analysis.

**Figure S5.** Comparative amino acid enrichment pathway scores on cells in the TME from day 21 and 28 post explanted B16-Ova melanoma tumor.

**Figure S6.** scRNA-seq analysis of expression of marker genes used to identify different cell types and signaling pathways in TME.

**Figure S7.** S1.0 remodulates metabolic pathways and enhances immune cell AA uptake.

### Supplementary Tables

**Table S1.** Biomarker genes used to identify divergent cell types

**Table S2.** Differentially expressed genes in TME treated with or without SPIKE 1.0 at 21 days post tumor cell implantation (PTCI)

- S1.0 vs Ctrl_ 21 days PTCI
- CAR-T vs Ctrl_ 21 days PTCI
- S1.0+CAR-T vs Ctrl_ 21 days PTCI

**Table S3.** Differentially expressed genes in TME treated with or without SPIKE 1.0 at 28 days PTCI

- S1.0 vs Ctrl_ 28 days PTCI
- CAR-T vs Ctrl_28 days PTCI
- S1.0+CAR-T vs Ctrl_28 days PTCI

**Table S4.** Metabolomics of tumours from treated tumor-bearing mice

**Table 5**. Primers used in this work

## References

1. Guo, C., Liu, J., and Zhang, Y. (2024). Current advances in bacteria-based cancer immunotherapy. European Journal of Immunology 54, 2350778.

2. Kwon, S.Y., Thi-Thu Ngo, H., Son, J., Hong, Y., and Min, J.J. (2024). Exploiting bacteria for cancer immunotherapy. Nat Rev Clin Oncol 21, 569–589. 10.1038/s41571-024-00908-9.

3. Gupta, K.H., Nowicki, C., Giurini, E.F., Marzo, A.L., and Zloza, A. (2021). Bacterial-Based Cancer Therapy (BBCT): Recent Advances, Current Challenges, and Future Prospects for Cancer Immunotherapy. Vaccines (Basel) 9. 10.3390/vaccines9121497.

4. Zhou, M., Tang, Y., Xu, W., Hao, X., Li, Y., Huang, S., Xiang, D., and Wu, J. (2023). Bacteria-based immunotherapy for cancer: a systematic review of preclinical studies. Front Immunol 14, 1140463. 10.3389/fimmu.2023.1140463.

5. Weeks, J.N., Galindo, C.L., Drake, K.L., Adams, G.L., Garner, H.R., and Ficht, T.A. (2010). Brucella melitensis VjbR and C12-HSL regulons: contributions of the N-dodecanoyl homoserine lactone signaling molecule and LuxR homologue VjbR to gene expression. BMC Microbiol 10, 167. 10.1186/1471-2180-10-167.

6. Guo, F., Das, J.K., Kobayashi, K.S., Qin, Q.M., T, A.F., Alaniz, R.C., Song, J., and Figueiredo, P. (2022). Live attenuated bacterium limits cancer resistance to CAR-T therapy by remodeling the tumor microenvironment. J Immunother Cancer 10. 10.1136/jitc-2021-003760.

7. Castano-Zubieta, M.R., Rossetti, C.A., Garcia-Gonzalez, D.G., Maurizio, E., Hensel, M.E., Rice-Ficht, A.C., Ficht, T.A., and Arenas-Gamboa, A.M. (2021). Evaluation of the safety profile of the vaccine candidate Brucella melitensis 16MDeltavjbR strain in goats. Vaccine 39, 617–625. 10.1016/j.vaccine.2020.11.033.

8. Arenas-Gamboa, A., Rice-Ficht, A., Fan, Y., Kahl-McDonagh, M., and Ficht, T. (2012). Extended safety and efficacy studies of the attenuated Brucella vaccine candidates 16MΔ vjbR and S19Δ vjbR in the immunocompromised IRF-1−/− mouse model. Clinical and Vaccine Immunology 19, 249–260.

9. Wang, Y., Bai, Y., Qu, Q., Xu, J., Chen, Y., Zhong, Z., Qiu, Y., Wang, T., Du, X., and Wang, Z. (2011). The 16MΔvjbR as an ideal live attenuated vaccine candidate for differentiation between Brucella vaccination and infection. Veterinary microbiology 151, 354–362.

10. Darbandi, A., Koupaei, M., Navidifar, T., Shahroodian, S., Heidary, M., and Talebi, M. (2022). Brucellosis control methods with an emphasis on vaccination: a systematic review. Expert Rev Anti Infect Ther 20, 1025–1035. 10.1080/14787210.2022.2066521.

11. Hensel, M.E., Garcia-Gonzalez, D.G., Chaki, S.P., Hartwig, A., Gordy, P.W., Bowen, R., Ficht, T.A., and Arenas-Gamboa, A.M. (2020). Vaccine Candidate Brucella melitensis 16MDeltavjbR Is Safe in a Pregnant Sheep Model and Confers Protection. mSphere 5. 10.1128/mSphere.00120-20.

12. Karimabad, M.N., Mahmoodi, M., Jafarzadeh, A., Darekordi, A., Hajizadeh, M.R., and Hassanshahi, G. (2019). Molecular Targets, Anti-cancer Properties and Potency of Synthetic Indole-3-carbinol Derivatives. Mini Rev Med Chem 19, 540–554. 10.2174/1389557518666181116120145.

13. Kaur, K., Verma, H., Gangwar, P., Jangid, K., Dhiman, M., Kumar, V., and Jaitak, V. (2024). Design, synthesis, in silico and biological evaluation of new indole based oxadiazole derivatives targeting estrogen receptor alpha. Bioorg Chem 147, 107341. 10.1016/j.bioorg.2024.107341.

14. Das, J.K., Guo, F., Hunt, C., Steinmeyer, S., Plocica, J.A., Kobayashi, K.S., Ding, Y., Jayaraman, A., Ficht, T.A., Alaniz, R.C., et al. (2022). A metabolically engineered bacterium controls autoimmunity and inflammation by remodeling the pro-inflammatory microenvironment. Gut Microbes 14, 2143222. 10.1080/19490976.2022.2143222.

15. Jia, D., Wang, Q., Qi, Y., Jiang, Y., He, J., Lin, Y., Sun, Y., Xu, J., Chen, W., Fan, L., et al. (2024). Microbial metabolite enhances immunotherapy efficacy by modulating T cell stemness in pan-cancer. Cell 187, 1651–1665 e1621. 10.1016/j.cell.2024.02.022.

16. Teymori, A., Mokhtari, S., Sedaghat, A., Mahboubi, A., and Kobarfard, F. (2023). Design, Synthesis, and Investigation of Cytotoxic Effects of 5-Hydroxyindole-3-Carboxylic Acid and Ester Derivatives as Potential Anti-breast Cancer Agents. Iran J Pharm Res 22, e133868. 10.5812/ijpr-133868.

17. Gurbatri, C.R., Lia, I., Vincent, R., Coker, C., Castro, S., Treuting, P.M., Hinchliffe, T.E., Arpaia, N., and Danino, T. (2020). Engineered probiotics for local tumor delivery of checkpoint blockade nanobodies. Sci Transl Med 12. 10.1126/scitranslmed.aax0876.

18. Wang, H., Xu, F., Yao, C., Dai, H., Xu, J., Wu, B., Tian, B., Shi, X., and Wang, C. (2024). Engineering bacteria for cancer immunotherapy by inhibiting IDO activity and reprogramming CD8+ T cell response. Proc Natl Acad Sci U S A 121, e2412070121. 10.1073/pnas.2412070121.

19. Yamazaki, Y., Kawano, Y., Yamanaka, A., and Maruyama, S. (2009). N– [(Dihydroxyphenyl)acyl]serotonins as potent inhibitors of tyrosinase from mouse and human melanoma cells. Bioorg Med Chem Lett 19, 4178–4182. 10.1016/j.bmcl.2009.05.115.

20. Granchi, C., Roy, S., Giacomelli, C., Macchia, M., Tuccinardi, T., Martinelli, A., Lanza, M., Betti, L., Giannaccini, G., Lucacchini, A., et al. (2011). Discovery of N-hydroxyindole-based inhibitors of human lactate dehydrogenase isoform A (LDH-A) as starvation agents against cancer cells. J Med Chem 54, 1599–1612. 10.1021/jm101007q.

21. Zhong, T., Sun, S., Zhao, M., Zhang, B., and Xiong, H. (2025). The mechanisms and clinical significance of CD8(+) T cell exhaustion in anti-tumor immunity. Cancer Biol Med 22, 460–480. 10.20892/j.issn.2095-3941.2024.0628.

22. Xie, Y.J., Dougan, M., Jailkhani, N., Ingram, J., Fang, T., Kummer, L., Momin, N., Pishesha, N., Rickelt, S., Hynes, R.O., and Ploegh, H. (2019). Nanobody-based CAR T cells that target the tumor microenvironment inhibit the growth of solid tumors in immunocompetent mice. Proc Natl Acad Sci U S A 116, 7624–7631. 10.1073/pnas.1817147116.

23. Ferrari, D.P., Ramos-Gomes, F., Alves, F., and Markus, M.A. (2024). KPC-luciferase-expressing cells elicit an anti-tumor immune response in a mouse model of pancreatic cancer. Sci Rep 14, 13602. 10.1038/s41598-024-64053-0.

24. Hildner, K., Edelson, B.T., Purtha, W.E., Diamond, M., Matsushita, H., Kohyama, M., Calderon, B., Schraml, B.U., Unanue, E.R., Diamond, M.S., et al. (2008). Batf3 deficiency reveals a critical role for CD8alpha+ dendritic cells in cytotoxic T cell immunity. Science 322, 1097–1100. 10.1126/science.1164206.

25. Spranger, S., Dai, D., Horton, B., and Gajewski, T.F. (2017). Tumor-Residing Batf3 Dendritic Cells Are Required for Effector T Cell Trafficking and Adoptive T Cell Therapy. Cancer Cell 31, 711–723 e714. 10.1016/j.ccell.2017.04.003.

26. Lapidot, T., Pflumio, F., Doedens, M., Murdoch, B., Williams, D.E., and Dick, J.E. (1992). Cytokine stimulation of multilineage hematopoiesis from immature human cells engrafted in SCID mice. Science 255, 1137–1141.

27. Shultz, L.D., Lyons, B.L., Burzenski, L.M., Gott, B., Chen, X., Chaleff, S., Kotb, M., Gillies, S.D., King, M., and Mangada, J. (2005). Human lymphoid and myeloid cell development in NOD/LtSz-scid IL2Rγnull mice engrafted with mobilized human hemopoietic stem cells. The Journal of Immunology 174, 6477–6489.

28. McCune, J., Kaneshima, H., Krowka, J., Namikawa, R., Outzen, H., Peault, B., Rabin, L., Shih, C.C., Yee, E., Lieberman, M., and, et al. (1991). The SCID-hu mouse: a small animal model for HIV infection and pathogenesis. Annu Rev Immunol 9, 399–429. 10.1146/annurev.iy.09.040191.002151.

29. Lopes, A., Bastiancich, C., Bausart, M., Ligot, S., Lambricht, L., Vanvarenberg, K., Ucakar, B., Gallez, B., Preat, V., and Vandermeulen, G. (2021). New generation of DNA-based immunotherapy induces a potent immune response and increases the survival in different tumor models. J Immunother Cancer 9. 10.1136/jitc-2020-001243.

30. Pappalardo, F., Martinez Forero, I., Pennisi, M., Palazon, A., Melero, I., and Motta, S. (2011). SimB16: modeling induced immune system response against B16-melanoma. PLoS One 6, e26523. 10.1371/journal.pone.0026523.

31. Hao, Y., Stuart, T., Kowalski, M.H., Choudhary, S., Hoffman, P., Hartman, A., Srivastava, A., Molla, G., Madad, S., Fernandez-Granda, C., and Satija, R. (2024). Dictionary learning for integrative, multimodal and scalable single-cell analysis. Nat Biotechnol 42, 293–304. 10.1038/s41587-023-01767-y.

32. Cibrian, D., and Sanchez-Madrid, F. (2017). CD69: from activation marker to metabolic gatekeeper. Eur J Immunol 47, 946–953. 10.1002/eji.201646837.

33. Sugiura, A., Beier, K.L., Chi, C., Heintzman, D.R., Ye, X., Wolf, M.M., Patterson, A.R., Cephus, J.Y., Hong, H.S., Lyssiotis, C.A., et al. (2023). Tissue-Specific Dependence of Th1 Cells on the Amino Acid Transporter SLC38A1 in Inflammation. bioRxiv. 10.1101/2023.09.13.557496.

34. Guo, C., You, Z., Shi, H., Sun, Y., Du, X., Palacios, G., Guy, C., Yuan, S., Chapman, N.M., Lim, S.A., et al. (2023). SLC38A2 and glutamine signalling in cDC1s dictate anti-tumour immunity. Nature 620, 200–208. 10.1038/s41586-023-06299-8.

35. Fridlender, Z.G., Sun, J., Kim, S., Kapoor, V., Cheng, G., Ling, L., Worthen, G.S., and Albelda, S.M. (2009). Polarization of tumor-associated neutrophil phenotype by TGF-beta: “N1” versus “N2” TAN. Cancer Cell 16, 183–194. 10.1016/j.ccr.2009.06.017.

36. Wang, X., Qiu, L., Li, Z., Wang, X.Y., and Yi, H. (2018). Understanding the Multifaceted Role of Neutrophils in Cancer and Autoimmune Diseases. Front Immunol 9, 2456. 10.3389/fimmu.2018.02456.

37. Chen, Q., Yin, H., Liu, S., Shoucair, S., Ding, N., Ji, Y., Zhang, J., Wang, D., Kuang, T., Xu, X., et al. (2022). Prognostic value of tumor-associated N1/N2 neutrophil plasticity in patients following radical resection of pancreas ductal adenocarcinoma. J Immunother Cancer 10. 10.1136/jitc-2022-005798.

38. Kerdkumthong, K., Nanarong, S., Roytrakul, S., Pitakpornpreecha, T., Tantimetta, P., Runsaeng, P., and Obchoei, S. (2024). Quantitative proteomics analysis reveals possible anticancer mechanisms of 5’-deoxy-5’-methylthioadenosine in cholangiocarcinoma cells. PLoS One 19, e0306060. 10.1371/journal.pone.0306060.

39. Li, Y., Wang, Y., and Wu, P. (2019). 5’-Methylthioadenosine and Cancer: old molecules, new understanding. J Cancer 10, 927–936. 10.7150/jca.27160.

40. Jin, J., Byun, J.K., Choi, Y.K., and Park, K.G. (2023). Targeting glutamine metabolism as a therapeutic strategy for cancer. Exp Mol Med 55, 706–715. 10.1038/s12276-023-00971-9.

41. Cheng, C.T., Qi, Y., Wang, Y.C., Chi, K.K., Chung, Y., Ouyang, C., Chen, Y.R., Oh, M.E., Sheng, X., Tang, Y., et al. (2018). Arginine starvation kills tumor cells through aspartate exhaustion and mitochondrial dysfunction. Commun Biol 1, 178. 10.1038/s42003-018-0178-4.

42. Grasmann, G., Smolle, E., Olschewski, H., and Leithner, K. (2019). Gluconeogenesis in cancer cells– repurposing of a starvation-induced metabolic pathway? Biochimica et Biophysica Acta (BBA)– Reviews on Cancer 1872, 24–36.

43. Kuo, M.T., Chen, H.H.W., Feun, L.G., and Savaraj, N. (2021). Targeting the Proline-Glutamine-Asparagine-Arginine Metabolic Axis in Amino Acid Starvation Cancer Therapy. Pharmaceuticals (Basel) 14. 10.3390/ph14010072.

44. Kober, C., Roewe, J., Schmees, N., Roese, L., Roehn, U., Bader, B., Stoeckigt, D., Prinz, F., Gorjanacz, M., Roider, H.G., et al. (2023). Targeting the aryl hydrocarbon receptor (AhR) with BAY 2416964: a selective small molecule inhibitor for cancer immunotherapy. J Immunother Cancer 11. 10.1136/jitc-2023-007495.

45. Ly, M., Rentas, S., Vujovic, A., Wong, N., Moreira, S., Xu, J., Holzapfel, N., Bhatia, S., Tran, D., Minden, M.D., et al. (2019). Diminished AHR Signaling Drives Human Acute Myeloid Leukemia Stem Cell Maintenance. Cancer Res 79, 5799–5811. 10.1158/0008-5472.CAN-19-0274.

46. Griffith, B.D., and Frankel, T.L. (2024). The Aryl Hydrocarbon Receptor: Impact on the Tumor Immune Microenvironment and Modulation as a Potential Therapy. Cancers (Basel) 16. 10.3390/cancers16030472.

47. Hezaveh, K., Shinde, R.S., Klotgen, A., Halaby, M.J., Lamorte, S., Ciudad, M.T., Quevedo, R., Neufeld, L., Liu, Z.Q., Jin, R., et al. (2022). Tryptophan-derived microbial metabolites activate the aryl hydrocarbon receptor in tumor-associated macrophages to suppress anti-tumor immunity. Immunity 55, 324–340 e328. 10.1016/j.immuni.2022.01.006.

48. Campesato, L.F., Budhu, S., Tchaicha, J., Weng, C.H., Gigoux, M., Cohen, I.J., Redmond, D., Mangarin, L., Pourpe, S., Liu, C., et al. (2020). Blockade of the AHR restricts a Treg-macrophage suppressive axis induced by L-Kynurenine. Nat Commun 11, 4011. 10.1038/s41467-020-17750-z.

49. Nayer, B., Tan, J.L., Alshoubaki, Y.K., Lu, Y.Z., Legrand, J.M.D., Lau, S., Hu, N., Park, A.J., Wang, X.N., Amann-Zalcenstein, D., et al. (2024). Local administration of regulatory T cells promotes tissue healing. Nat Commun 15, 7863. 10.1038/s41467-024-51353-2.

50. Shi, H., Chapman, N.M., Wen, J., Guy, C., Long, L., Dhungana, Y., Rankin, S., Pelletier, S., Vogel, P., Wang, H., et al. (2019). Amino Acids License Kinase mTORC1 Activity and Treg Cell Function via Small G Proteins Rag and Rheb. Immunity 51, 1012–1027 e1017. 10.1016/j.immuni.2019.10.001.

51. Varanasi, S.K., Chen, D., Liu, Y., Johnson, M.A., Miller, C.M., Ganguly, S., Lande, K., LaPorta, M.A., Hoffmann, F.A., Mann, T.H., et al. (2025). Bile acid synthesis impedes tumor-specific T cell responses during liver cancer. Science 387, 192–201. 10.1126/science.adl4100.

52. Geiger, R., Rieckmann, J.C., Wolf, T., Basso, C., Feng, Y., Fuhrer, T., Kogadeeva, M., Picotti, P., Meissner, F., Mann, M., et al. (2016). L-Arginine Modulates T Cell Metabolism and Enhances Survival and Anti-tumor Activity. Cell 167, 829–842 e813. 10.1016/j.cell.2016.09.031.

53. Yao, C.C., Sun, R.M., Yang, Y., Zhou, H.Y., Meng, Z.W., Chi, R., Xia, L.L., Ji, P., Chen, Y.Y., Zhang, G.Q., et al. (2023). Accumulation of branched-chain amino acids reprograms glucose metabolism in CD8(+) T cells with enhanced effector function and anti-tumor response. Cell Rep 42, 112186. 10.1016/j.celrep.2023.112186.

54. Zheng, Y., Yao, Y., Ge, T., Ge, S., Jia, R., Song, X., and Zhuang, A. (2023). Amino acid metabolism reprogramming: shedding new light on T cell anti-tumor immunity. J Exp Clin Cancer Res 42, 291. 10.1186/s13046-023-02845-4.

55. Park, J., Hsueh, P.C., Li, Z., and Ho, P.C. (2023). Microenvironment-driven metabolic adaptations guiding CD8(+) T cell anti-tumor immunity. Immunity 56, 32–42. 10.1016/j.immuni.2022.12.008.

56. Ganjoo, S., Gupta, P., Corbali, H.I., Nanez, S., Riad, T.S., Duong, L.K., Barsoumian, H.B., Masrorpour, F., Jiang, H., Welsh, J.W., and Cortez, M.A. (2023). The role of tumor metabolism in modulating T-Cell activity and in optimizing immunotherapy. Front Immunol 14, 1172931. 10.3389/fimmu.2023.1172931.

57. Craze, M.L., El-Ansari, R., Aleskandarany, M.A., Cheng, K.W., Alfarsi, L., Masisi, B., Diez-Rodriguez, M., Nolan, C.C., Ellis, I.O., Rakha, E.A., and Green, A.R. (2019). Glutamate dehydrogenase (GLUD1) expression in breast cancer. Breast Cancer Res Treat 174, 79–91. 10.1007/s10549-018-5060-z.

58. Xu, W., Patel, C.H., Zhao, L., Sun, I.H., Oh, M.H., Sun, I.M., Helms, R.S., Wen, J., and Powell, J.D. (2023). GOT1 regulates CD8(+) effector and memory T cell generation. Cell Rep 42, 111987. 10.1016/j.celrep.2022.111987.

59. Colpitts, S.L., Dalton, N.M., and Scott, P. (2009). IL-7 receptor expression provides the potential for long-term survival of both CD62Lhigh central memory T cells and Th1 effector cells during Leishmania major infection. J Immunol 182, 5702–5711. 10.4049/jimmunol.0803450.

60. Akkari, L., Amit, I., Bronte, V., Fridlender, Z.G., Gabrilovich, D.I., Ginhoux, F., Hedrick, C.C., and Ostrand-Rosenberg, S. (2024). Defining myeloid-derived suppressor cells. Nat Rev Immunol 24, 850– 857. 10.1038/s41577-024-01062-0.

61. Khan, A.U., Melzer, F., Sayour, A.E., Shell, W.S., Linde, J., Abdel-Glil, M., El-Soally, S., Elschner, M.C., Sayour, H.E.M., Ramadan, E.S., et al. (2021). Whole-Genome Sequencing for Tracing the Genetic Diversity of Brucella abortus and Brucella melitensis Isolated from Livestock in Egypt. Pathogens 10. 10.3390/pathogens10060759.

62. Bankhead, P., Loughrey, M.B., Fernandez, J.A., Dombrowski, Y., McArt, D.G., Dunne, P.D., McQuaid, S., Gray, R.T., Murray, L.J., Coleman, H.G., et al. (2017). QuPath: Open source software for digital pathology image analysis. Sci Rep 7, 16878. 10.1038/s41598-017-17204-5.

63. Hao, Y., Hao, S., Andersen-Nissen, E., Mauck, W.M., 3rd, Zheng, S., Butler, A., Lee, M.J., Wilk, A.J., Darby, C., Zager, M., et al. (2021). Integrated analysis of multimodal single-cell data. Cell 184, 3573–3587 e3529. 10.1016/j.cell.2021.04.048.

64. Franzen, O., Gan, L.M., and Bjorkegren, J.L.M. (2019). PanglaoDB: a web server for exploration of mouse and human single-cell RNA sequencing data. Database (Oxford) 2019. 10.1093/database/baz046.

65. Aibar, S., Gonzalez-Blas, C.B., Moerman, T., Huynh-Thu, V.A., Imrichova, H., Hulselmans, G., Rambow, F., Marine, J.C., Geurts, P., Aerts, J., et al. (2017). SCENIC: single-cell regulatory network inference and clustering. Nat Methods 14, 1083–1086. 10.1038/nmeth.4463.

66. Cabello, A.L., Wells, K., Peng, W., Feng, H.Q., Wang, J., Meyer, D.F., Noroy, C., Zhao, E.S., Zhang, H., Li, X., et al. (2024). Brucella-driven host N-glycome remodeling controls infection. Cell Host Microbe 32, 588–605 e589. 10.1016/j.chom.2024.03.003.

67. Wells, K.M., He, K., Pandey, A., Cabello, A., Zhang, D., Yang, J., Gomez, G., Liu, Y., Chang, H., Li, X., et al. (2022). Brucella activates the host RIDD pathway to subvert BLOS1-directed immune defense. Elife 11. 10.7554/eLife.73625.

68. Qin, Q.M., Pei, J., Ancona, V., Shaw, B.D., Ficht, T.A., and de Figueiredo, P. (2008). RNAi screen of endoplasmic reticulum-associated host factors reveals a role for IRE1alpha in supporting Brucella replication. PLoS Pathog 4, e1000110. 10.1371/journal.ppat.1000110.

69. Fernandez-Prada, C.M., Zelazowska, E.B., Nikolich, M., Hadfield, T.L., Roop, R.M., 2nd, Robertson, G.L., and Hoover, D.L. (2003). Interactions between Brucella melitensis and human phagocytes: bacterial surface O-Polysaccharide inhibits phagocytosis, bacterial killing, and subsequent host cell apoptosis. Infect Immun 71, 2110–2119. 10.1128/IAI.71.4.2110-2119.2003.

70. Yen, K.M., and Karl, M.R. (1992). Identification of a new gene, tmoF, in the Pseudomonas mendocina KR1 gene cluster encoding toluene-4-monooxygenase. J. Bacteriol. 174, 7253–7261. 10.1128/jb.174.22.7253-7261.1992.

71. Fleetwood, A.J., Lee, M.K.S., Singleton, W., Achuthan, A., Lee, M.C., O’Brien-Simpson, N.M., Cook, A.D., Murphy, A.J., Dashper, S.G., Reynolds, E.C., and Hamilton, J.A. (2017). Metabolic Remodeling, Inflammasome Activation, and Pyroptosis in Macrophages Stimulated by Porphyromonas gingivalis and Its Outer Membrane Vesicles. Front Cell Infect Microbiol 7, 351. 10.3389/fcimb.2017.00351.

72. Weischenfeldt, J., and Porse, B. (2008). Bone Marrow-Derived Macrophages (BMM): Isolation and Applications. CSH Protoc 2008, pdb prot5080. 10.1101/pdb.prot5080.

73. Waclawikova, B., Bullock, A., Schwalbe, M., Aranzamendi, C., Nelemans, S.A., van Dijk, G., and El Aidy, S. (2021). Gut bacteria-derived 5-hydroxyindole is a potent stimulant of intestinal motility via its action on L-type calcium channels. PLoS Biol 19, e3001070. 10.1371/journal.pbio.3001070.

74. Gaikwad, S.R., Nguyen, H.M., Srivastava, S.K., and Wood, L. (2023). Abstract LB201: A novel combination of Listeria-based immunotherapy and repurposed drug combats colorectal cancer. Cancer Research 83, LB201–LB201.

75. Ren, Y., Kumar, A., Das, J.K., Peng, H.Y., Wang, L., Balllard, D., Xiong, X., Ren, X., Zhang, Y., Yang, J.M., and Song, J. (2022). Tumorous expression of NAC1 restrains antitumor immunity through the LDHA-mediated immune evasion. J Immunother Cancer 10. 10.1136/jitc-2022-004856.

76. Das, J.K., Ren, Y., Kumar, A., Peng, H.Y., Wang, L., Xiong, X., Alaniz, R.C., de Figueiredo, P., Ren, X., Liu, X., et al. (2022). Elongation factor-2 kinase is a critical determinant of the fate and antitumor immunity of CD8(+) T cells. Sci Adv 8, eabl9783. 10.1126/sciadv.abl9783.

77. Rosenberg, S.A., and Restifo, N.P. (2015). Adoptive cell transfer as personalized immunotherapy for human cancer. Science 348, 62–68. 10.1126/science.aaa4967.

78. Ma, Y., Li, J., Wang, H., Chiu, Y., Kingsley, C.V., Fry, D., Delaney, S.N., Wei, S.C., Zhang, J., Maitra, A., and Yee, C. (2020). Combination of PD-1 Inhibitor and OX40 Agonist Induces Tumor Rejection and Immune Memory in Mouse Models of Pancreatic Cancer. Gastroenterology 159, 306–319 e312. 10.1053/j.gastro.2020.03.018.

79. Suarez-Esquivel, M., Chaves-Olarte, E., Moreno, E., and Guzman-Verri, C. (2020). Brucella Genomics: Macro and Micro Evolution. Int J Mol Sci 21. 10.3390/ijms21207749.

80. Krypotou, E., Townsend, G.E., Gao, X., Tachiyama, S., Liu, J., Pokorzynski, N.D., Goodman, A.L., and Groisman, E.A. (2023). Bacteria require phase separation for fitness in the mammalian gut. Science 379, 1149–1156.

81. Valenzano, G., Russell, S.N., Go, S., O’Neill, E., and Jones, K.I. (2024). Using Spectral Flow Cytometry to Characterize Anti-Tumor Immunity in Orthotopic and Subcutaneous Mouse Models of Cancer. Curr Protoc 4, e70032. 10.1002/cpz1.70032.

82. Nagarajan, A., Scoggin, K., Gupta, J., Aminian, M., Adams, L.G., Kirby, M., Threadgill, D., and Andrews-Polymenis, H. (2024). Collaborative Cross mice have diverse phenotypic responses to infection with Methicillin-resistant Staphylococcus aureus USA300. PLoS Genet 20, e1011229. 10.1371/journal.pgen.1011229.

83. Blank, H.M., Hammer, S.E., Boatright, L., Roberts, C., Heyden, K.E., Nagarajan, A., Tsuchiya, M., Brun, M., Johnson, C.D., Stover, P.J., et al. (2024). Late-life dietary folate restriction reduces biosynthesis without compromising healthspan in mice. Life Sci Alliance 7. 10.26508/lsa.202402868.

84. Osorio, D., and Cai, J.J. (2021). Systematic determination of the mitochondrial proportion in human and mice tissues for single-cell RNA-sequencing data quality control. Bioinformatics 37, 963–967. 10.1093/bioinformatics/btaa751.

85. Kuleshov, M.V., Jones, M.R., Rouillard, A.D., Fernandez, N.F., Duan, Q., Wang, Z., Koplev, S., Jenkins, S.L., Jagodnik, K.M., Lachmann, A., et al. (2016). Enrichr: a comprehensive gene set enrichment analysis web server 2016 update. Nucleic Acids Res 44, W90–97. 10.1093/nar/gkw377.

86. Zhang, L., and Nishi, H. (2022). Transcriptome analysis of Homo sapiens and Mus musculus reveals mechanisms of CD8+ T cell exhaustion caused by different factors. PLoS One 17, e0274494. 10.1371/journal.pone.0274494.

87. Aluganti Narasimhulu, C., and Singla, D.K. (2021). Amelioration of diabetes-induced inflammation mediated pyroptosis, sarcopenia, and adverse muscle remodelling by bone morphogenetic protein-7. J Cachexia Sarcopenia Muscle 12, 403–420. 10.1002/jcsm.12662.

88. Salazar, C., Armenta, J.M., and Shulaev, V. (2012). An UPLC-ESI-MS/MS Assay Using 6-Aminoquinolyl-N-Hydroxysuccinimidyl Carbamate Derivatization for Targeted Amino Acid Analysis: Application to Screening of Arabidopsis thaliana Mutants. Metabolites 2, 398–428. 10.3390/metabo2030398.

89. Apetoh, L., Quintana, F.J., Pot, C., Joller, N., Xiao, S., Kumar, D., Burns, E.J., Sherr, D.H., Weiner, H.L., and Kuchroo, V.K. (2010). The aryl hydrocarbon receptor interacts with c-Maf to promote the differentiation of type 1 regulatory T cells induced by IL-27. Nat Immunol 11, 854–861. 10.1038/ni.1912.

