## Supplemental Figures 1 to 7 for "A Synthetic Microbial Therapy Rewires Antitumor Immunity Across Multiple Cancer Types"

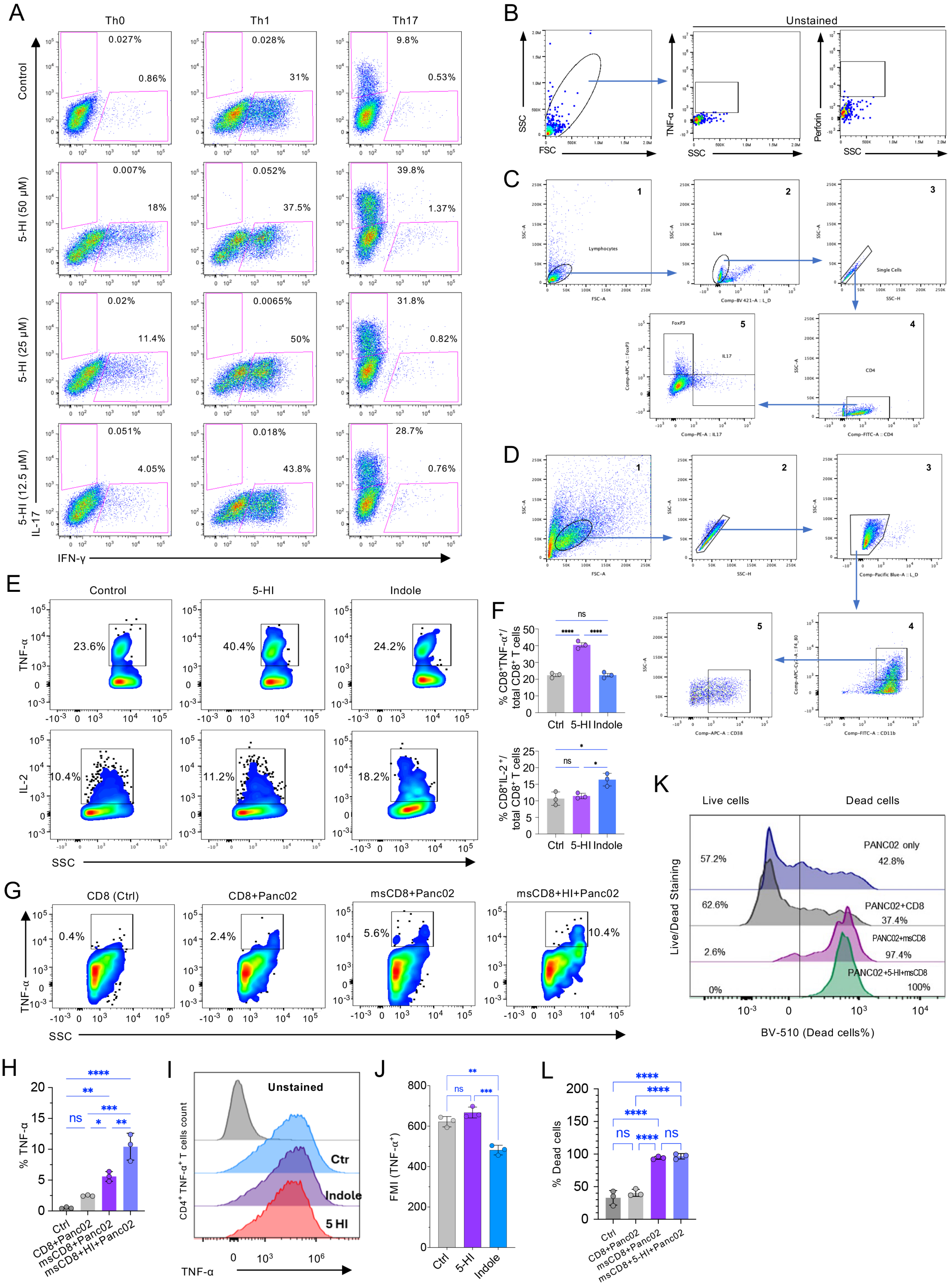
A
Th0
Th1
Th17
Control
5-HI (50 μM)
5-HI (25 μM)
5-HI (12.5 μM)
IL-17
IFN-γ
0.027%
0.86%
0.028%
31%
9.8%
0.53%
0.007%
18%
0.052%
37.5%
39.8%
1.37%
0.02%
11.4%
0.0065%
50%
31.8%
0.82%
0.051%
4.05%
0.018%
43.8%
28.7%
0.76%
B
Unstained
SSC
FSC
TNF-α
Perforin
C
1
2
3
4
5
Lymphocytes
Live
Single Cells
FoxP3
IL17
CD4
D
E
Control
5-HI
Indole
TNF-α
IL-2
SSC
23.6%
40.4%
24.2%
10.4%
11.2%
18.2%
F
% CD8+TNF-α+/ total CD8+ T cells
% CD8+IL-2+/ total CD8+ T cells
ns
****
*
Ctrl
5-HI
Indole
G
CD8 (Ctrl)
CD8+Panc02
msCD8+Panc02
msCD8+HI+Panc02
0.4%
2.4%
5.6%
10.4%
K
Live cells
Dead cells
Live/Dead Staining
PANC02 only
57.2%
42.8%
PANC02+CD8
62.6%
37.4%
PANC02+msCD8
2.6%
97.4%
PANC02+5-HI+msCD8
0%
100%
BV-510 (Dead cells%)
H
% TNF-α
Ctrl
CD8+Panc02
msCD8+Panc02
msCD8+HI+Panc02
I
Unstained
Ctr
Indole
5 HI
CD4+ TNF-α+ T cells count
TNF-α
J
FMI (TNF-α+)
Ctrl
5-HI
Indole
L
% Dead cells
Ctrl
CD8+Panc02
msCD8+Panc02
msCD8+5-HI+Panc02

**Figure S1. Hydroxyindole (HI) increases cytokine production and cytotoxicity of CD8$^+$ T cells, related to Figure 1**

(A) 5-HI induces Th1 and Th17 responses in CD4 T-cells. CD4$^+$CD25$^-$ naïve T-cells were sorted (>98% purity) from naïve B6 mouse spleens, seeded at 2.0 x $10^5$ per well in plates coated with anti-CD3 and anti-CD28 agonist antibodies under Th0 (IL-2 plus no skew cytokines), Th1 (IL-2 plus IL-12 cytokine), or Th17 (TGFb plus IL-6) conditions in the presence of physiologic concentrations of microbiota-derived metabolites HI (50, 25, 12.5 uM final), or not (Control). After 3 days, cells were restimulated with PMA and Ionomycin (5 hours) and stained for CD4, and for IFN- γ and IL-17 to indicate activated Th1 or Th17 T-cells.

(B-D) Representative flow cytometry gating strategies. (B) Gating strategy for flow cytometry analysis of Figure 1A. Cells were first identified based on forward scatter (FSC) and side scatter (SSC) parameters to exclude debris and select the primary cell population. The unstained plot for TNF-α and Perforin is shown in the Figure. (C) Gating strategy for flow cytometry analysis shown in Figure 1E. 1) Lymphocytes were gated (SSC-A vs FSC-A). 2) Dead cells were removed from the analysis using Zombie Violet™. 3) Doublets were removed from living cells (Live/Dead$^-$) using FSC-A and FSC-H. 4) Singlet live Lymphocyte gate was further analyzed for CD4 T cells population, and 5) CD4$^+$ T cells selected for further characterization of FoxP3$^+$ and IL17$^+$ Tregs and Th17 cells respectively. (D) Gating strategy for flow cytometry analysis of Figure 1C. 1) Bone marrow derived macrophages (BMDM) were gated (SSC-A vs FSC-A). 2) Doublets were removed from BMDM pool using FSC-A and FSC-H. 3) Dead cells were removed from the analysis using Zombie Violet™. 4) Live BMDM gate was further analyzed for CD11b and F4_80 to to gate live macrophages. 5) and then CD11b$^+$ CF4/80$^+$ cells selected for further expression of CD38$^+$ M1 macrophages.

(E) 5-HI regulates cytokines TNF-α and IL-2 expression levels in human CD8$^+$ T cells. Human CD8$^+$ T cells were isolated from human peripheral blood mononuclear cells (PBMCs) and activated with anti-human CD3/CD28 antibodies.

(F) Quantification of the indicated protein marker-positive in CD8$^+$ T cell populations.

(G, H) The CFSE$^+$ CD8$^+$ T cells were analyzed for intracellular cytokine production TNF-α determined by flowcytometric analysis (G) and quantitation of TNF-α$^+$CD8$^+$ cell population (H).

(I, J) TNF-α production in CD4$^+$ T cells after 5-HI treatment. CD4$^+$ T cells isolated from C57BL/6 mice were activated with anti-CD3/CD28 antibodies and treated with 5-HI as described the Methods section, TNF-α production in CD4$^+$ T cells was determined by flowcytometric analysis (I) and quantification of TNF-α production in the CD4$^+$ T cells (J).

(K, L) Flow-cytometry analysis of cells for dead-live staining using Aqua zombie dead live staining dye (K) and quantitation of dead cells (%) recovered from flowcytometric analysis (L). The representative data are shown from a ratio of 1:1 (CD8$^+$ T cells to cancer cells). msCD8: Meso-expressing CFSE$^+$ CD8$^+$ T cells.

Data represent mean ± standard error of measurement (SEM) from at least three independent experiments. *, $p<0.05$; **, $p<0.01$; ***, $p<0.001$; ****, $p<0.0001$.

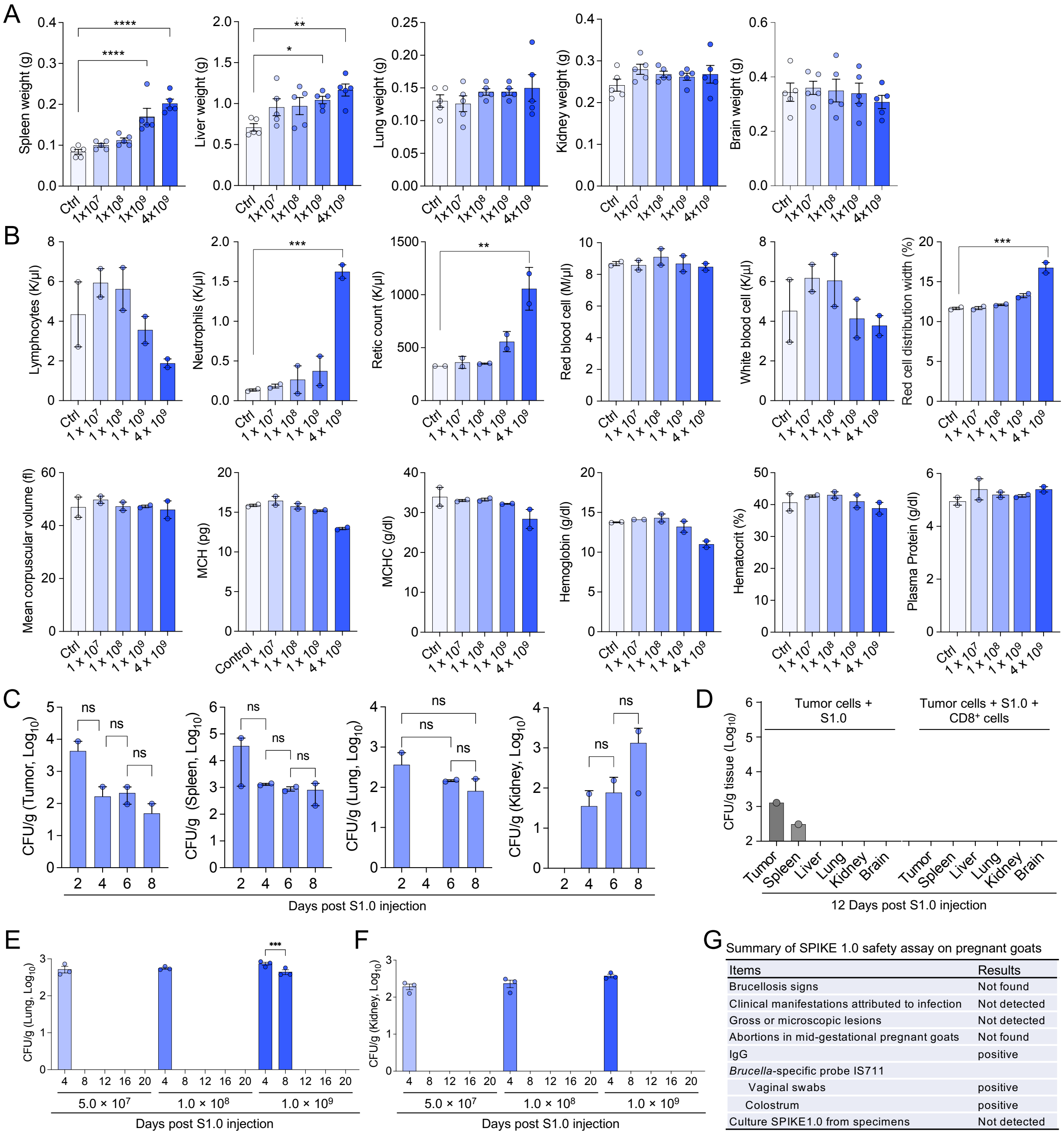


Summary of SPIKE 1.0 safety assay on pregnant goats

| Items | Results |
|---|---|
| Brucellosis signs | Not found |
| Clinical manifestations attributed to infection | Not detected |
| Gross or microscopic lesions | Not detected |
| Abortions in mid-gestational pregnant goats | Not found |
| IgG | positive |
| *Brucella*-specific probe IS711 | |
| Vaginal swabs | positive |
| Colostrum | positive |
| Culture SPIKE1.0 from specimens | Not detected |

**Figure S2. Safety assessment of SPIKE 1.0 (S1.0) in mice and pregnant goat models, related to Figure 1**

(A) Organ weight analysis of mice (n = 5/group) injected with different doses of S1.0 at 14 days post injection (DPI).

(B) CBC analysis of blood contents in mice (n = 5/group) administered with different bacterial doses of S1.0 at 14 DPI. MCH: mean corpuscular hemoglobin, MCHC: mean corpuscular hemoglobin concentration. Data represent mean ± SEM from two independent experiments (effective samples: n = 2, 3 in experiment 1 and 2, respectively).

(C) Colony-forming unit (CFU) analysis of S1.0 distribution in tumor, spleen, lung, and kidney of B16-Ova melanoma tumor-bearing mice administrated with the dose of $5.0\times10^7$ bacteria of S1.0.

(D-F) CFU analysis of S1.0 distribution in tumor and the indicated organs in Lewis lung carcinoma (LLC1) tumor-bearing mice administrated with S1.0 ($5.0\times10^7$) and with or without adoptively transferred $CD8^+$ T-cells (D), or in lung (E) and kidney (F) of the LLC1 tumor-bearing mice treated with the indicated doses of S1.0 alone. n = 5 mice/group. Data represent mean ± SEM from at least two independent experiments for panel C to F.

(G) Summary of safety analysis of S1.0 in pregnant goats. n = 7/treatment.

*: $p \leq 0.05$, **: $p \leq 0.01$, ***: $p \leq 0.001$, and ****: $p \leq 0.0001$.

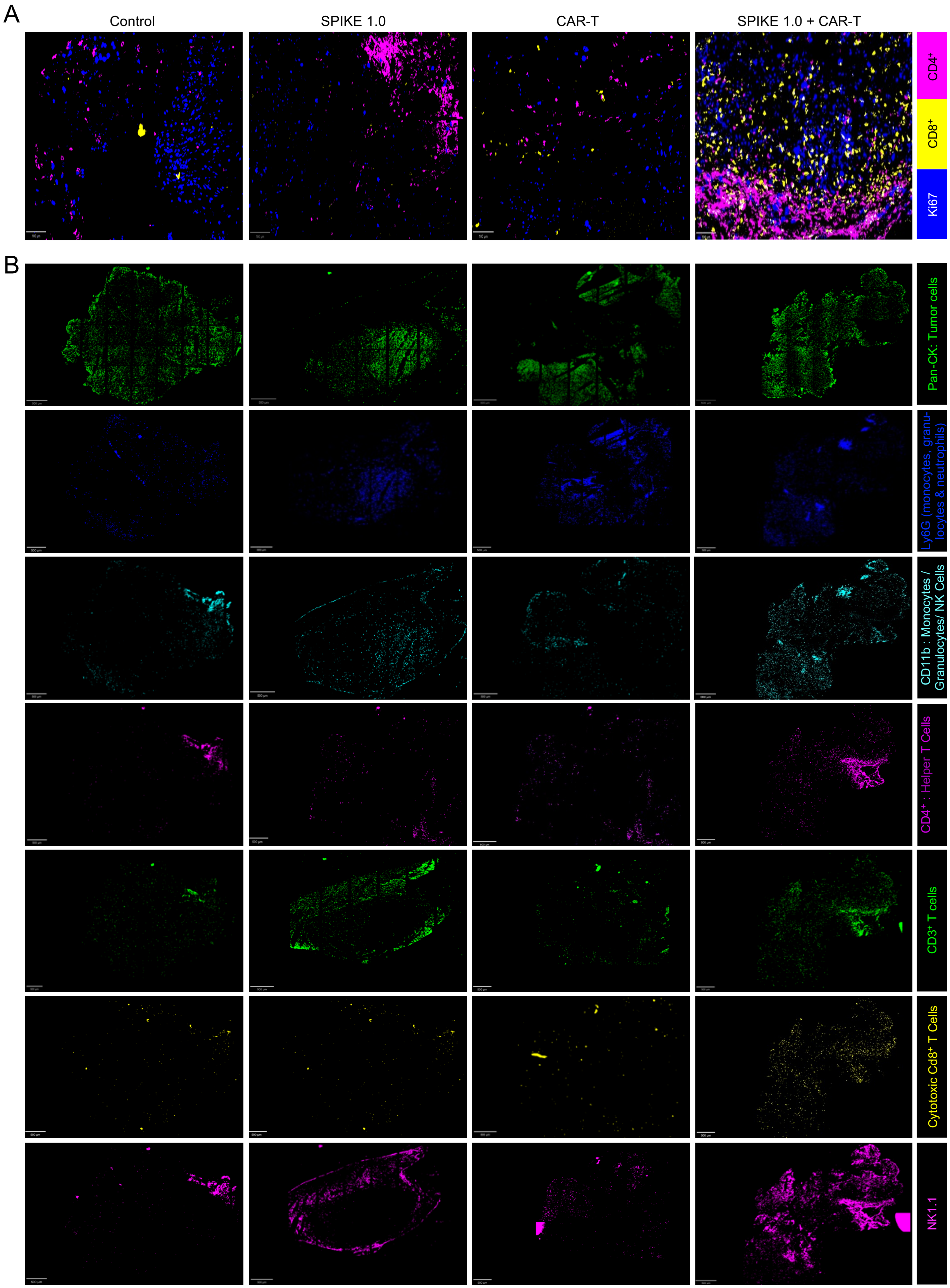
A
Control
SPIKE 1.0
CAR-T
SPIKE 1.0 + CAR-T
CD4+
CD8+
Ki67
B
Pan-CK: Tumor cells
Ly6G (monocytes, granu-locytes & neutrophils)
CD11b : Monocytes / Granulocytes/ NK Cells
CD4+ : Helper T Cells
CD3+ T cells
Cytotoxic Cd8+ T Cells
NK1.1

1

**Figure S3. Chip-cytometry multiplex imaging analysis of biomarker expression in the TME of B16-Ova melanoma tumor-bearing mice at day 28 PTCI, related to Figure 4**

(A) Representative multiparametric images showing $CD4^+$ T cells, CD8+ T cells infiltration and Ki-67 proliferation into the TME. Bar: 100 µm.

(B) Representative single channel of chip-cytometry multiplex images showing different immune cells in the TME. Bars: 500 µm.

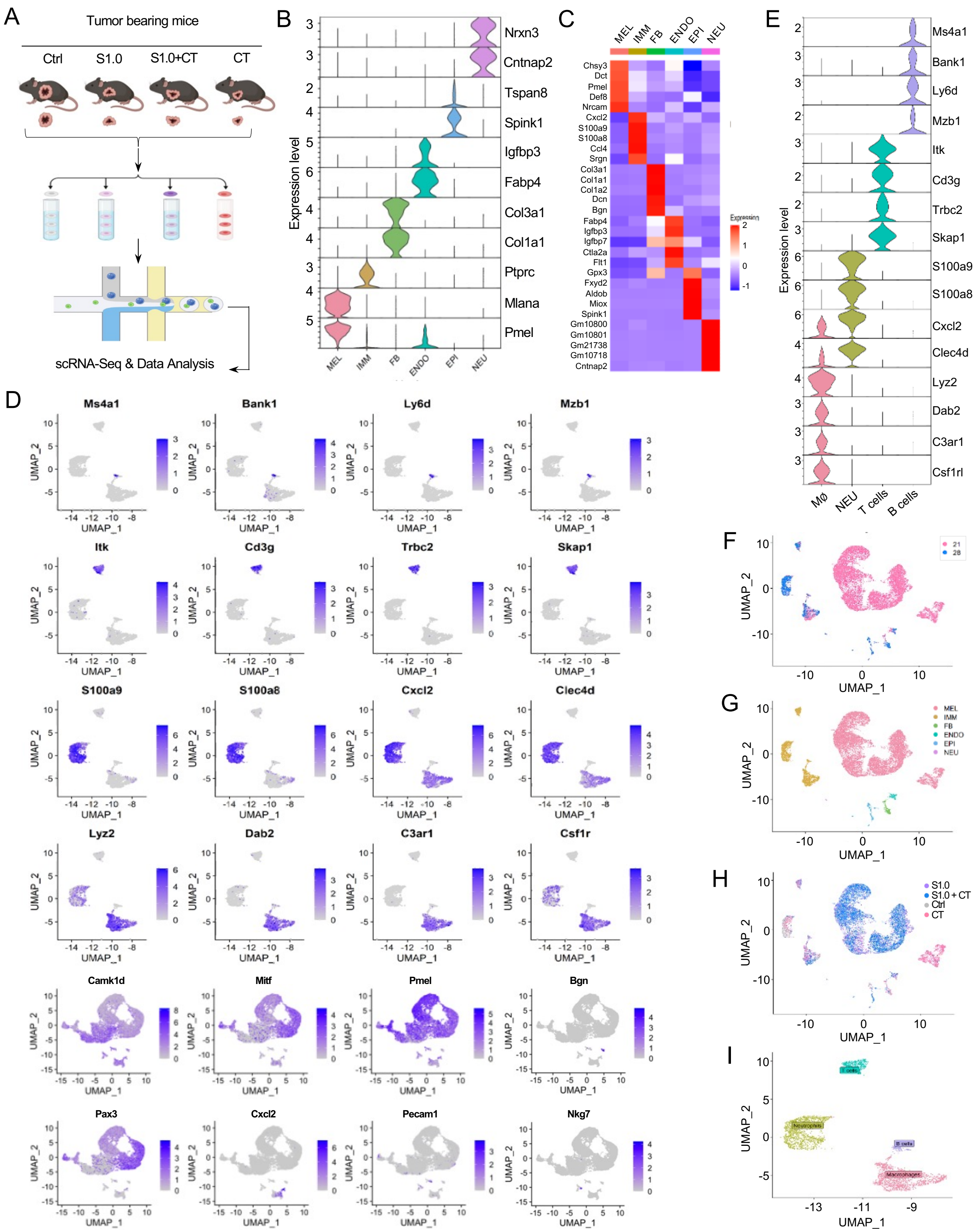


1 **Figure S4. Identification of different cell types by scRNA-seq analysis, related to Figure 6**

(A) Schematic of B16-Ova melanoma tumor sample preparation and scRNA-seq experimental approach.

(B) Representative violin plots of different gene markers used to identify the indicated cell types. MEL: melanocytes, IMM: immune cells, FB: fibroblasts, ENDO: endothelial cells, EPI: epithelial cells, and NEU: neutrophils.

(C) Heatmap showing the expression of different gene markers used to identify MEL, IMM, FB, ENDO, EP, and NEU.

(D) UMAP clustering analysis of the different gene markers used for the identification of macrophages, neutrophils, T cells and B cells.

(E) Violin plots of expression of the indicated gene markers used to identify macrophages, neutrophils, T cells and B cells.

(F) Seurat based UMAP clustering of all different cell types derived from TME on Day 21 and Day 28 PTCI.

(G) UMAP clustering and classification of the different cell types (MEL, IMM, FB, ENDO, EPI, and NEU) in scRNA-seq analysis at Day 21 and Day 28 PTCI.

(H) UMAP clustering and representation of the different cell types at Day 21 and Day 28 PTCI based on the treatment strategies.

(I) Seurat based UMAP clustering of immune cell types (B cells, T cells, and Neutrophils) in TME at day 28 PTCI. n = 3.

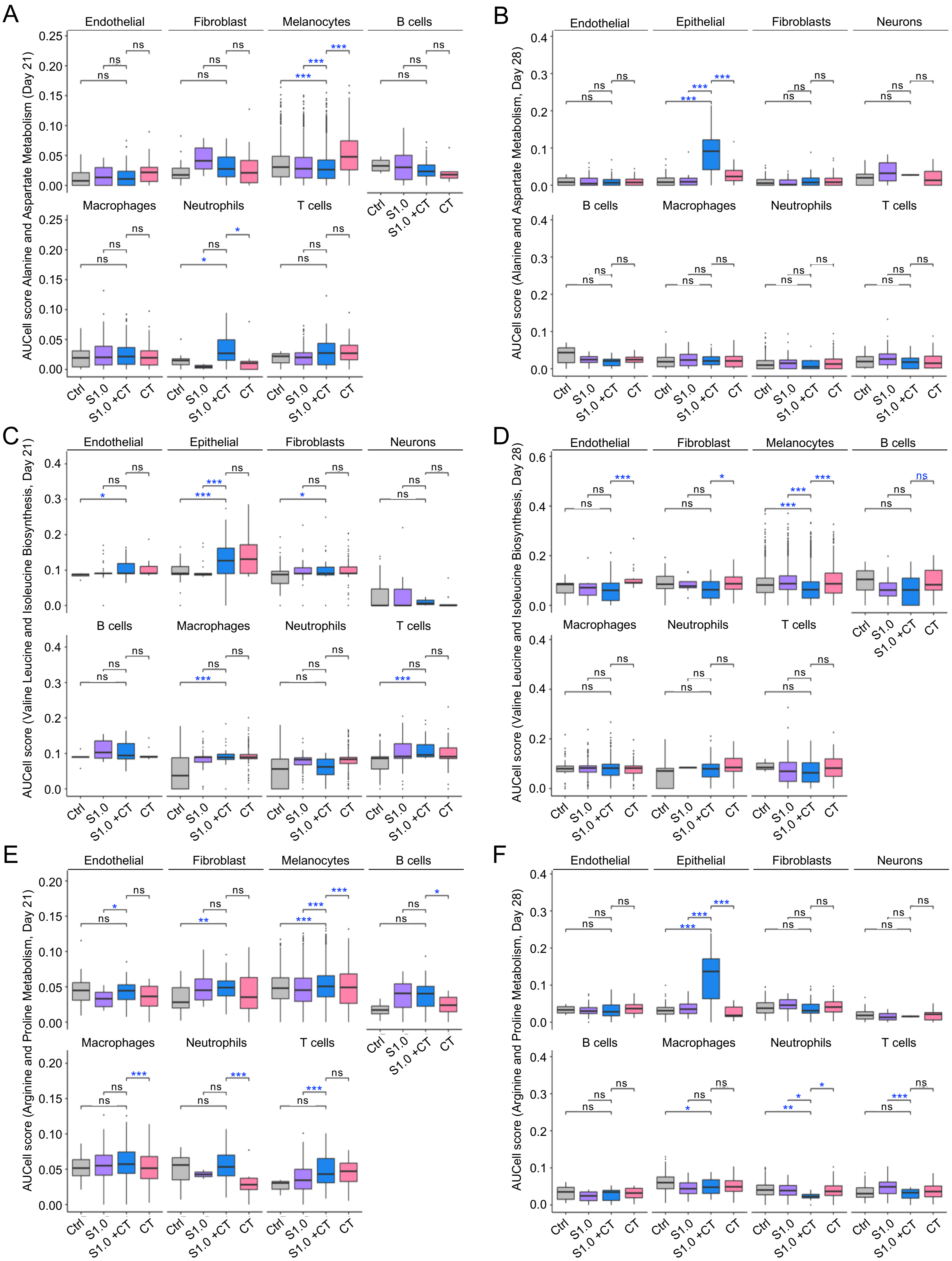
A
Endothelial
Fibroblast
Melanocytes
B cells
Macrophages
Neutrophils
T cells
AUCell score Alanine and Aspartate Metabolism (Day 21)
Ctrl
S1.0
S1.0 +CT
CT
B
Endothelial
Epithelial
Fibroblasts
Neurons
B cells
Macrophages
Neutrophils
T cells
AUCell score (Alanine and Aspartate Metabolism, Day 28)
C
Endothelial
Epithelial
Fibroblasts
Neurons
B cells
Macrophages
Neutrophils
T cells
AUCell score (Valine Leucine and Isoleucine Biosynthesis, Day 21)
D
Endothelial
Fibroblast
Melanocytes
B cells
Macrophages
Neutrophils
T cells
AUCell score (Valine Leucine and Isoleucine Biosynthesis, Day 28)
E
Endothelial
Fibroblast
Melanocytes
B cells
Macrophages
Neutrophils
T cells
AUCell score (Arginine and Proline Metabolism, Day 21)
F
Endothelial
Epithelial
Fibroblasts
Neurons
B cells
Macrophages
Neutrophils
T cells
AUCell score (Arginine and Proline Metabolism, Day 28)

**Figure S5. Comparative amino acid enrichment pathway scores on cells in the TME from day 21 and 28 post explanted B16-Ova melanoma tumor, related to Figure 6**

(A-B) Alanine and aspartate metabolism pathway at day 21 (A) and day 28 (B) PTCI.

(C-D) Valine, leucine and isoleucine biosynthesis pathway at day 21 (C) and day 28 (D) PTCI.

(E-F) Arginine and proline metabolism at day 21 (E) and day 28 (F) PTCI.

n = 3. *: $p < 0.05$, **: 0.01, ***: 0.001, ****: 0.0001. ns: not significant.

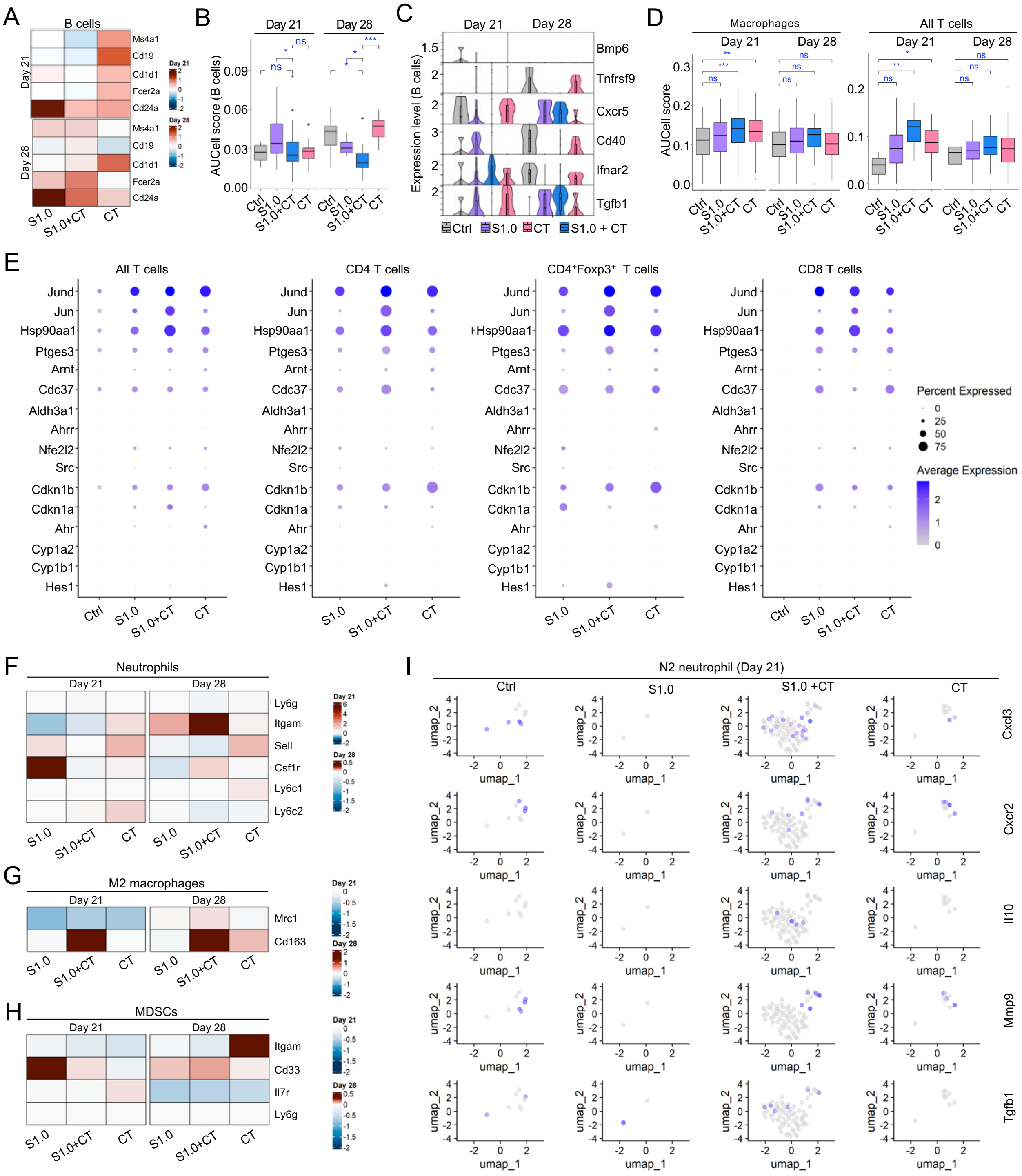
A
B cells
Day 21
Day 28
Ms4a1
Cd19
Cd1d1
Fcer2a
Cd24a
S1.0
S1.0+CT
CT
B
Day 21
Day 28
AUCell score (B cells)
0.09
0.06
0.03
0.00
ns
*
***
Ctrl
S1.0
S1.0+CT
CT
C
Day 21
Day 28
Expression level (B cells)
Bmp6
Tnfrsf9
Cxcr5
Cd40
Ifnar2
Tgfb1
Ctrl
S1.0
CT
S1.0 + CT
D
Macrophages
All T cells
Day 21
Day 28
AUCell score
**
***
ns
E
All T cells
CD4 T cells
CD4⁺Foxp3⁺ T cells
CD8 T cells
Jund
Jun
Hsp90aa1
Ptges3
Arnt
Cdc37
Aldh3a1
Ahrr
Nfe2l2
Src
Cdkn1b
Cdkn1a
Ahr
Cyp1a2
Cyp1b1
Hes1
Percent Expressed
0
25
50
75
Average Expression
2
1
0
F
Neutrophils
Day 21
Day 28
Ly6g
Itgam
Sell
Csf1r
Ly6c1
Ly6c2
G
M2 macrophages
Mrc1
Cd163
H
MDSCs
Itgam
Cd33
Il7r
Ly6g
I
N2 neutrophil (Day 21)
Ctrl
S1.0
S1.0 +CT
CT
umap_1
umap_2
Cxcl3
Cxcr2
Il10
Mmp9
Tgfb1

1

**Figure S6. scRNA-seq analysis of expression of marker genes used to identify different cell types and signaling pathways in TME, related to Figure 6**

(A) Expression analysis of marker genes that identify B cells in TME with the indicated treatments at day 21 and 28 PTCI.

(B) AUCell score analysis of cytokine-cytokine receptor signaling pathway in B cells.

(C) Violin plots showing the differential expressed genes of the cytokine-cytokine receptor signaling pathway in B cells among different groups.

(D) AUCell analysis of gene sets of the aryl hydrocarbon receptor (AhR) signaling pathway in macrophages and in all T cells.

(E) Expression analysis of the marker genes in the AhR signaling pathways in the indicated T cells.

(F-H) Expression analysis of marker genes that identify neutrophils (F), M2 macrophages (G), and myeloid-derived suppressor cells (MDSCs) (H) in TME at day 21 and 28 PTCI.

(I) UMAP clustering analysis showing the expression of marker genes that identify N2 neutrophil in TME at day 21 PTCI.

n = 3. *: $p < 0.05$, **: 0.01, ***: 0.001. ns: not significant.

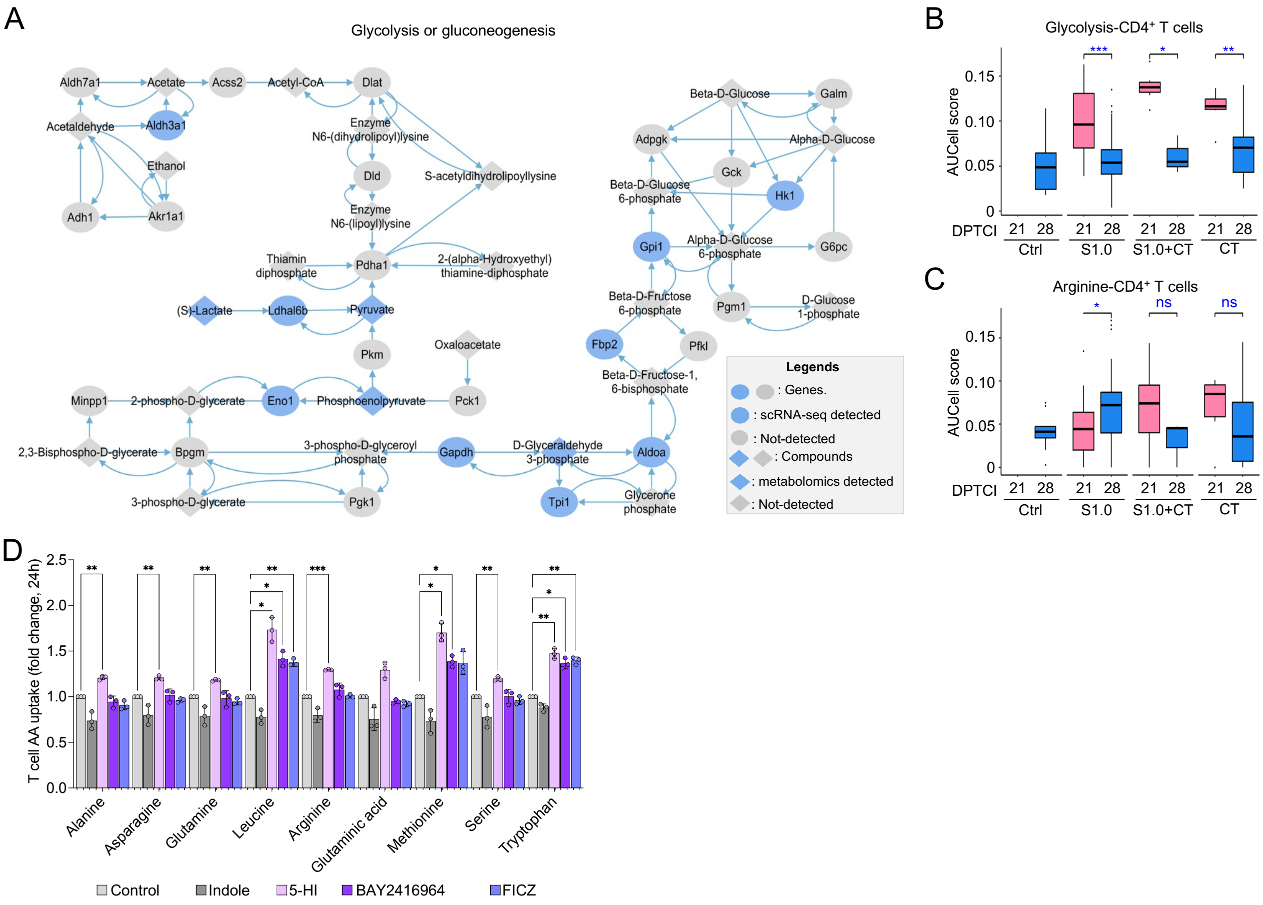


**Figure S7. S1.0 remodulates metabolic pathways and enhances immune cell AA uptake, related to Figure 7**

(A) Network map analysis of glycolysis or gluconeogenesis pathway in the TEM of B16-Ova tumor treated with Ctrl, SC or S1.0 at Day 21. The analyzed datasets are derived from the combined scRNA-seq data (Table S2-S3) and metabolomics (Table S4).

(B, C) Comparison of glycolysis levels (B) and arginine biosynthesis levels (C) in CD4$^+$ T cells in the TME of B16-Ova tumor across different treatments at 21 and 28 days PTCI. The analyzed dataset is from scRNA-seq data (Table S2-S3).

(D) T cell AA uptake profiling analysis at 24 h post co-culture with 5-HI and other indicated small molecules. T cells were isolated from mouse spleen. FICZ: 6-formylindolo [3, 2-b] carbazole. n=3. *, **, ***: $p < 0.05$, 0.01, and 0.001, respectively.
