## Supplemental Table 1-5, and will be used for the link to file on the preprint sites for "A Synthetic Microbial Therapy Rewires Antitumor Immunity Across Multiple Cancer Types": Das et al_SPIKE 1.0 Table S1.docx

**Table S1.** Biomarker genes used to identify divergent cell types

| Cell Type | Marker Genes | Purpose |
| --- | --- | --- |
| Melanocytes | Pmel, Mlana, Pax3, Dct | Identify melanocytes in TME |
| Immune cells | Ptprc, Cxcl2, S100a9, S100a8, Lyz2 | Identify Immune in TME |
| Endothelial Cells | Fabp4, Cd93, Pecam1, Igfbp7, Adgrf5, Ptprb | Identify endothelial cells in TME |
| Epithelial Cells | Krt18, Tspan8, Krt7 | Identify epithelial cells in TME |
| Fibroblasts | Col3a1, Col1a1, Col1a2, Dcn, Bgn, Fbn1 | Identify Fibroblasts in TME |
| Neurons | Nrnx3, Rbfox1, Nrg3 | Identify Neurons in TME |
| immune cell types |  |  |
| Macrophages (M1, M2) | Lyz2, Apoe, Csf1r, Cd68 | Identify Macrophages in TME |
| Neutrophils (N1, N2) | Csf3r, S100a8, S100a9, Lcn2 | Identify Neutrophils in TME |
| T cells | Itk, Skap1, Cd3g, Trbc2 | Identify T cells in TME |
| B cells | Ms4a1, Cd79a, Cd79b, Bank1 | Identify B cells in TME |
