## Supplemental Table 1-5, and will be used for the link to file on the preprint sites for "A Synthetic Microbial Therapy Rewires Antitumor Immunity Across Multiple Cancer Types": Das et al_SPIKE 1.0 Table S5.docx

**Table S5.** Primers used in this work

| **Gene/**  **sequence** | **Forward primer** | **Reverse primer** | **Purpose** |
| --- | --- | --- | --- |
| Slc38a10 | CCACCTATGACAGCCTGGAC | GCCCACTCGGATCATCTCTG | qPCR validation |
| Malt-1 | GAACTGAGCGACTTCCTACAGG | AACTGTCCAGCCAACACTGCCT | qPCR validation |
| Slc1a5 | CTGCCTGTGAAGGACATCTCCT | CTCGGCATCTTGGTTCGATCCA | qPCR validation |
| Adora2a | CACGCAGAGTTCCATCTTCAGC | CCCAGCAAATCGCAATGATGCC | qPCR validation |
| Gls | CAGAAGGCACAGACATGGTTGG | CAAGGTGGCAGCCATCACACTT | qPCR validation |
| GLUT-1 | CTTCATTGTGGGCATGTGCTTC | AGGTTCGGCCTTTGGTCTCAG | qPCR validation |
| GoT-1 | TGCTACTGGGATGCGGAGAAGA | TGCATGACAGCAGCGATCTGCT | qPCR validation |
| β-actin | CATTGCTGACAGGATGCAGAAGG | TGCTGGAAGGTGGACAGTGAGG | qPCR validation |
| IS711 | ATTCAATCTGATGGCGTTCC | GCCTATGATGCCGATCACTT | SPIKE detection |
| TMO | CGGCCTGATAAATTGAGGAA | GCAACTTTCTCCGCTACCTG | SPIKE target gene detection |
| IS711-Probe: CACTGGAACGTGTTGGATTG (/56-**FAM**/CACTGGAAC/ZEN/GTGTTGGATTG/3IABkFQ/) | | | Fluorescent dye attachment and fluorescence detection |
| TMO-Probe: AATGTCGGCATTTCCAGTTC (/5**Cy5**/AATGTCGGC/TAO/ATTTCCAGTTC/3IAbRQSp/) | | | Fluorescent dye attachment and fluorescence detection |
